# Integration of machine learning and pan-genomics expands the biosynthetic landscape of RiPP natural products

**DOI:** 10.1101/2020.05.19.104752

**Authors:** Alexander M. Kloosterman, Peter Cimermancic, Somayah S. Elsayed, Chao Du, Michalis Hadjithomas, Mohamed S. Donia, Michael A. Fischbach, Gilles P. van Wezel, Marnix H. Medema

## Abstract

Most clinical drugs are based on microbial natural products, with compound classes including polyketides (PKS), non-ribosomal peptides (NRPS), fluoroquinones and ribosomally synthesized and post-translationally modified peptides (RiPPs). While variants of biosynthetic gene clusters (BGCs) for known classes of natural products are easy to identify in genome sequences, BGCs for new compound classes escape attention. In particular, evidence is accumulating that for RiPPs, subclasses known thus far may only represent the tip of an iceberg. Here, we present decRiPPter (Data-driven Exploratory Class-independent RiPP TrackER), a RiPP genome mining algorithm aimed at the discovery of novel RiPP classes. DecRiPPter combines a Support Vector Machine (SVM) that identifies candidate RiPP precursors with pan-genomic analyses to identify which of these are encoded within operon-like structures that are part of the accessory genome of a genus. Subsequently, it prioritizes such regions based on the presence of new enzymology and based on patterns of gene cluster and precursor peptide conservation across species. We then applied decRiPPter to mine 1,295 *Streptomyces* genomes, which led to the identification of 42 new candidate RiPP families that could not be found by existing programs. One of these was studied further and elucidated as a novel subfamily of lanthipeptides, designated Class V. Two previously unidentified modifying enzymes are proposed to create the hallmark lanthionine bridges. Taken together, our work highlights how novel natural product families can be discovered by methods going beyond sequence similarity searches to integrate multiple pathway discovery criteria.

**Code and data availability:** The source code of DecRiPPter is freely available online at https://github.com/Alexamk/decRiPPter. Results of the data analysis are available online at http://www.bioinformatics.nl/~medem005/decRiPPter_strict/index.html and http://www.bioinformatics.nl/~medem005/decRiPPter_mild/index.html (for the strict and mild filters, respectively). All training data and code used to generate these, as well as outputs of the data analyses, are available on Zenodo at <u>doi:10.5281/zenodo.3834818</u>.

## Introduction

The introduction of antibiotics in the 20^th^ century contributed hugely to extend the human life span. However, the increase in antibiotic resistance and the concomitant steep decline in the number of new compounds discovered via high-throughput screening^1,2^, means that we again face huge challenges to treat infections by multi-drug resistant bacteria^3^. The low return of investment of high throughput screening is due to dereplication, in other words, the rediscovery of bioactive compounds that have been identified before^4,5^. A revolution in our understanding was brought about by the development of next-generation sequencing technologies. Actinobacteria are the most prolific producers of bioactive compounds, including some two-thirds of the clinical antibiotics^6,7^. Mining of the genome sequences of these bacteria revealed a huge repository of previously unseen biosynthetic gene clusters (BGCs), highlighting that their potential as producers of bioactive molecules had been grossly underestimated^6,8,9^. However, these BGCs are often not expressed under laboratory conditions, most likely because the environmental cues that activate their expression in their original habitat are missing^10,11^. To circumvent these issues, a common strategy is to select a candidate BGC and force its expression by expression of the pathway-specific activator or via expression of the BGC in a heterologous host^12^. However, these methods are time-consuming, while it is hard to predict the novelty and utility of the compounds they produce.

To improve the success of genome mining-based drug discovery, many bioinformatic tools have been developed for identification and prioritization of BGCs. These tools often rely on conserved genetic markers present in BGCs of certain natural products, such as polyketides (PKS), non-ribosomal peptide synthetases (NRPS) and terpenes^13–15^. While these methods have unearthed vast amounts of uncharacterized BGCs, they further expand on previously characterized classes of natural products. This raises the question of whether entirely novel classes of natural products could still be discovered. A few genome mining methods, such as ClusterFinder^16^ and EvoMining^17,18^, have tried to tackle this problem. These methods either use criteria true of all BGCs or build around the evolutionary properties of gene families found in BGCs, rather than using specific BGC-class-specific genetic markers. While the lack of clear genetic markers may result in a higher number of false positives, these methods have indeed charted previously uncovered biochemical space and led to the discovery of new natural products.

One class of natural products whose expansion has been fueled by the increased amount of genomic sequences available is that of the ribosomally synthesized and post-translationally modified peptides (RiPPs)^19^. RiPPs are characterized by a unifying biosynthetic theme: a small gene encodes a short precursor peptide, which is extensively modified by a series of enzymes that typically recognize the N-terminal part of the precursor called the leader peptide, and finally cleaved to yield the mature product^20^. Despite this common biosynthetic logic, RiPP modifications are highly diverse. The latest comprehensive review categorizes RiPPs into roughly 20 different classes^19^, such as lanthipeptides, lasso peptides and thiopeptides. Each of these classes is characterized by one or more specific modifications, such as the thioether bridge in lanthipeptides or the knot-like structure of lasso peptides. Despite the extensive list of known classes and modifications, new RiPP classes are still being found. Newly identified RiPP classes often carry unusual modifications, such as D-amino acids^21^, addition of unnatural amino acids^22,23^, β-amino acids^24^, or new variants of thioether crosslinks^25,26^. These discoveries strongly indicate that the RiPP genomic landscape remains far from completely charted, and that novel types of RiPPs with new and unique biological activities may yet be uncovered. However, RiPPs pose a unique and major challenge to genome-based pathway identification attempts: unlike in the case of NRPSs and PKSs, there are no universally conserved enzyme families or enzymatic domains that are found across all RiPP pathways. Rather, each class of RiPPs comprises its own unique set of enzyme families to post-translationally modify the precursor peptides belonging to that class. Hence, while biosynthetic gene clusters (BGCs) for known RiPP classes can be identified using conventional genome mining algorithms, a much more elaborate strategy is required to automate the identification of novel RiPP classes.

Several methods have made progress in tackling this challenge. ‘Bait-based’ approaches such as RODEO^27,28^ and RiPPer^29^ identify RiPP BGCs by looking for homologues of RiPP tailoring enzymes (RTEs) of interest, and facilitate identifying these RTEs in novel contexts to find many new RiPP BGCs. However, these methods still require a known query RTE from a known RiPP subclass. Another tool recently described, NeuRiPP, is capable of predicting precursors independent of RiPP subclass, but is limited to precursor analysis^30^. Yet another tool, DeepRiPP, can detect novel RiPP BGCs that are chemically far removed from known examples, but is mainly designed to identify new members of known classes^31^. In the end, an algorithm for the discovery of BGCs encoding novel RiPP classes will need to integrate various sources of information to reliably identify genomic regions that are likely to encode RiPP precursors along with previously undiscovered RTEs.

Here, we present decRiPPter (Data-driven Exploratory Class-independent RiPP TrackER), an integrative algorithm for the discovery of novel classes of RiPPs, without requiring prior knowledge of their specific modifications or core enzymatic machinery. DecRiPPter employs a Support Vector Machine (SVM) classifier that predicts RiPP precursors regardless of RiPP subclass, and combines this with pan-genomic analysis to identify which putative precursor genes are located within specialized genomic regions that encode multiple enzymes and are part of the accessory genome of a genus. Sequence similarity networking of the resulting precursors and gene clusters then facilitates further prioritization. Applying this method to the gifted natural product producer genus *Streptomyces*, we identified 42 new RiPP family candidates. Experimental characterization of a widely distributed candidate RiPP BGC led to the discovery of a novel lanthipeptide that was produced by a previously unknown enzymatic machinery.

## Results

### RiPP BGC discovery by detection of genomic islands with characteristics typical of RiPP BGCs

Given the promise of RiPPs as a source for novel natural products, we set out to construct a platform to facilitate identification of novel RiPP classes. Since no criteria could be used that are specific for individual RiPP classes, we used three criteria that generally apply to RiPP BGCs: 1) they contain one or more ORFs for a precursor peptide; 2) they contain genes encoding modifying machinery in an operon-like gene cluster together with precursor gene(s); 3) they have a sparse distribution within the wider taxonomic group in which they are found. To focus on novel RiPP classes, we added a fourth criterion: 4) they have no direct similarity to BGCs of known classes (Figure 1).

**Figure 1.**
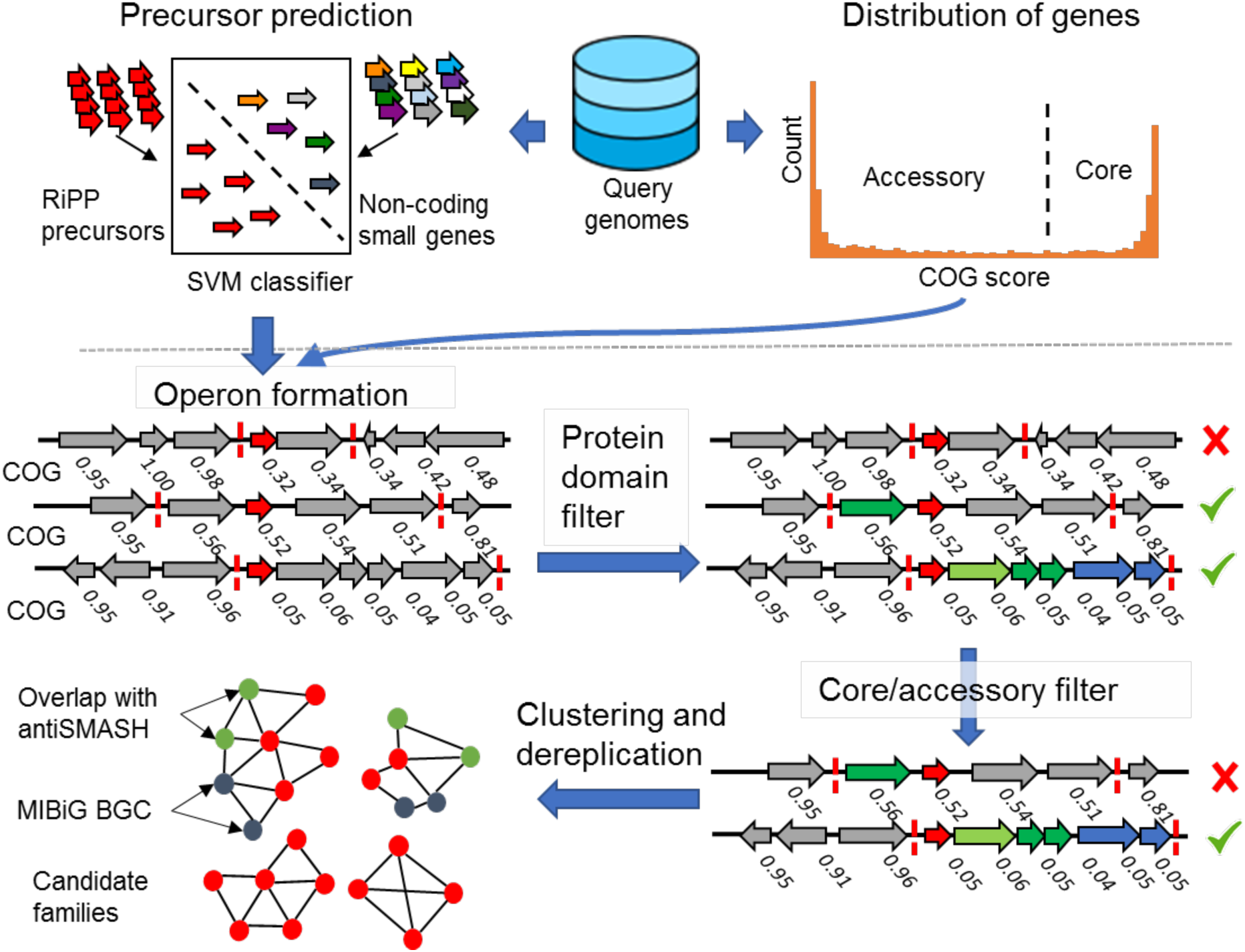
decRiPPter pipeline for the detection of novel RiPP families. From a given group of genomes, all genes smaller than 100 amino acids are analyzed by the SVM classifier, which finds candidate precursors. The gene clusters formed around the precursors are analyzed for specific protein domains. In addition, all COG scores are calculated to act as an additional filter, and to aid in gene cluster detection. The remaining gene clusters are clustered together and with MIBiG gene clusters to dereplicate and organize the results. In addition, overlap with antiSMASH detected BGCs is analyzed (**4**).

For the first criterion, we trained an SVM classifier to distinguish between RiPP precursors and other peptides. A collection of 175 known RiPP precursors, gathered from RiPP clusters from the MIBiG repository^32^ was used as a positive training set. For the negative training set, we generated a set of 20,000 short non-precursor sequences, consisting of 10,000 randomly selected short proteins (<175 amino acids long) from Uniprot without measurable similarity to RiPP precursors (representative of gene encoding proteins but not RiPP precursors), and 10,000 translated intergenic sequences between a stop codon and the next start codon of sizes 30-300 nt taken from 10 genomes across the bacterial tree of life (representative of spurious ORFs that do not encode proteins). From both positive and negative training set sequences, 36 different features were extracted describing the amino acid composition and physicochemical properties of the protein/peptide sequences, as well as localized enrichment of amino acids prone to modification by RTEs. Based on these, a support vector machine was trained (see details in Methods section). To make sure that this classifier could predict precursors independent of RiPP subclass, we trained it on all possible subsets of the positive training set in which one of the RiPP subclasses was entirely left out (a strategy we termed leave-one-class-out cross-validation). Typically, the classifier was still capable of predicting the class that was left out, with an area-under-receiver operating characteristics curve of 0.955.

For the second criterion, we made use of the fact that the majority of RiPP BGCs appear to contain the genes encoding the precursor and the core biosynthetic enzymes in the same strand orientation within close intergenic distance (81.6% of MIBiG RiPPs). Therefore, candidate gene clusters are formed from the genes that appear to reside in an operon with predicted precursor genes, based on intergenic distance and the COG scores calculated (see description of third criterion below, the Methods section and Figure S1). These gene clusters are then analyzed for protein domains that could constitute the modifying machinery (Figure 1b). Rather than restricting ourselves to specific protein domains, we constructed a broad dataset of Pfam and TIGRFAM domains that are linked to an E.C. number using InterPro mappings^33^. This dataset was extended with a previously curated set of Pfam domains found to be prevalent in the positive training set of the ClusterFinder algorithm^34^, and manually curated, resulting in a set of 4,131 protein domains. We also constructed Pfam^35^ and TIGRFAM^36^ domain datasets of transporters, regulators and peptidases, as well as a dataset consisting of known RiPP modifying domains to provide more detailed annotation and allow specific filtering of RiPP BGCs based on the presence of each of these types of Pfam domains (Supplemental Document 2).

For the third criterion, we sought to distinguish specialized genomic regions from conserved genomic regions. Indeed, most BGCs are sparingly distributed among genomes, with even closely related strains showing differences in their BGC repertoires^37–39^. We therefore developed an algorithm that separates the ‘core’ genome from the ‘accessory’ genome, by comparing all genes in a group of query genomes from the same taxon (typically a genus), and identifying the frequency of occurrence of each gene within that group of genomes (Figures 1c and S2). For the purpose of comparing genes between genomes, we reasoned that it was more straightforward to identify groups of functionally closely related genes that also include recent paralogues, due to the complexities of dealing with orthology relationships across large numbers of genomes (especially for biosynthetic genes that are known to have a discontinuous taxonomic distribution and may undergo frequent duplications^40^).

Therefore, decRiPPter first identifies the distribution of sequence identity values of protein-coding genes that can confidently be assigned to be orthologues, and uses this distribution to find groups of genes across genomes with orthologue-like mutual similarity. To identify a set of high-confidence orthologues, decRiPPter looks for genomic loci between which at least three contiguous genes are each other’s bidirectional best hits (BBHs, using DIAMOND^41^) between all possible genome pairs of the group of genomes analyzed, and assigned the center genes of these loci orthologue status, termed a true conserved orthologous gene (trueCOG)^42^. Since many orthologues are missed by only considering orthologues based on BBHs^43^, and to also include recent paralogues, we then further expanded the list of homologues with orthologue-like similarity by dynamically determining a cutoff between each genome pair based on the similarity of the trueCOGs shared between those genomes. This cutoff is used to find all highly similar gene pairs, which are then clustered with the Markov Clustering Algorithm (MCL^44^) into ‘clusters of orthologous genes’ (COGs). The number of COG members found for each gene is divided by the number of genomes in the query to get a COG score ranging from 0 to 1, reflecting how widespread the gene is across the set of query genomes. To validate our calculations, we analyzed the COG-scores of the highly conserved single-copy BUSCO gene set from OrthoDB^45,46^, as well as the COG-scores of the genes in the gene clusters predicted by antiSMASH. In line with our expectations, homologs of the BUSCO gene set averaged COG-scores of 0.95 (Figure S5), while the COG-scores of the antiSMASH gene clusters were much lower, averaging 0.311 +- 0.249 for all BGCs, and 0.234 +- 0.166 for RiPP BGCs (Figure S6). While the COG-scoring method requires a group of genomes to be analyzed rather than a single genome, we believe that the extra calculation significantly contributes in filtering false positives (see Table 1 and Figure S4). In addition, the COG scores aid in the gene cluster identification based on the assumption that gene clusters are generally sets of genes with similar absence/presence patterns across species (see Methods section).

**Table 1.** Increasing the strictness of the filter used on the found gene clusters results in a higher saturation of RiPP BGCs.

| Filter | Filter details | Number of detected gene clusters | Number of detected gene clusters overlapping with antiSMASH RiPP BGCs | Percentage of detected gene clusters overlapping with RiPP BGCs |
| --- | --- | --- | --- | --- |
| None | - | 718268 | 5908 | 0.8% |
| Mild | Gene cluster COG score: $\leq 0.25$<br>In the gene cluster:<br>$\geq 3$ genes<br>$\geq 2$ biosynthetic genes<br>In or around the gene cluster:<br>$\geq 1$ transporter | 21419 | 1678 | 7.8% |
| Strict | Gene cluster COG score: $\leq 0.10$<br>In the gene cluster:<br>$\geq 3$ genes<br>$\geq 2$ biosynthetic genes<br>In or around the gene cluster:<br>$\geq 1$ transporter<br>$\geq 1$ regulator<br>$\geq 1$ peptidase | 2471 | 357 | 14.4% |

For the final criterion, the algorithm dereplicates the identified clusters by comparing them to known RiPP BGCs. All putative BGCs are clustered based on domain content and precursor similarity using sequence similarity networking^47^, and compared to known RiPP BGCs from MIBiG^48^. In addition, the overlap between predicted RiPP BGCs and gene clusters found by antiSMASH^49^ is determined (Figure 1).

### decRiPPter identifies 42 candidate novel RiPP classes in *Streptomyces*

While RiPPs are found in many different microorganisms, their presence in streptomycetes reflects perhaps the most diverse array of RiPP classes within a single genus. Streptomycete*s* produce a broad spectrum of RiPPs, namely lanthipeptides^50^, lasso peptides^27^, linear azol(in)e-containing peptides (LAPs)^51^, thiopeptides^52^, thioamide-containing peptides^29^ and bottromycins^53^. Their potential as RiPP producers is further highlighted by a recent study showcasing the diversity of lanthipeptide BGCs in *Streptomyces* and other actinobacteria^54^. Given the large variety of different families of natural products produced by this genus, we hypothesized it to be a likely source of novel RiPP classes, and sought to exhaustively mine it.

We started by running the pipeline described above on all publicly available *Streptomyces* genomes (1,295 genomes) from NCBI (Supplemental Document 3). Due to computational limits, the genomes were split into ten randomly selected groups to calculate the frequency of distribution of each gene (COG-scores). In general, the number of genomes that could be grouped together and the resulting cutoffs were found to vary with the amount of minimum trueCOGs required (Figure S3A). To make sure that as many genomes as possible could be compared at once, we set the cutoff for minimum number of trueCOGs at 10. Despite the low cutoff, the distribution of similarity scores between genome pairs still resembled a Gaussian distribution (Figure S3B). The bimodal distribution of the resulting COG-scores showed that the majority of the genes were either conserved in only a small portion of the genomes, or present in almost all genomes (Figure S4).

We then scanned all predicted products of genes as well as predicted ORFs in intergenic regions shorter than 100 amino acids (total 7.19 * 10^7^) with the SVM classifier. While by far most of the queries scored below 0.5, a peak of queries scoring from 0.9 to 1.0 was observed (Figure S7). Seeking to be inclusive at this stage, we set the cutoff at 0.9, resulting in 1.32*10^6^ candidate precursors passing this initial filter, thus filtering out 98.2 % of all candidates. Eliminating candidate precursors whose genes were completely overlapping reduced the number to 8.17*10^5^ precursors (1.1 %). While, most probably, the vast majority of these are not RiPP precursors, it provides a suitably sized set of candidates to then enter the next stages of the decRiPPter workflow.

In our analyses, we found that the majority of RiPP BGCs contain the majority of biosynthetic genes on the same strand orientation as the precursor (MIBiG: 81.6%; antiSMASH RiPP BGCs: 73.1%). We therefore formed gene clusters using only the genes on the same strand as the predicted precursor. To create a training set, we divided all known RiPP BGCs and all antiSMASH RiPP BGCs found in the analyzed genome sequences into sections where each section contained only genes on the same strand. The core section was defined as the section that contained the most biosynthetic genes as detected by antiSMASH or as annotated in the MIBiG database. These sections were used as training sets to finetune distance and COG cutoffs for our gene cluster methods.

In a simple gene cluster method, genes were joined only using the intergenic distances as a cutoff. Using this method, we found that at a distance of 750 nucleotides, all MIBiG core sections were covered, and 91% of all antiSMASH core sections (Figure S8AB). However, using only distance may cause the gene cluster formation to overshoot into regions not associated with the BGC (e.g. Figure S2). We therefore created an alternative method, called the ‘island method’. In this method, each gene is first joined with immediately adjacent genes that lie in the same strand orientation and have very small intergenic regions (<=50 nucleotides), to form islands. These islands may subsequently be combined if they have similar average COG-scores (see materials and methods). We found that with this method, we could confidently cover our validation set, while slightly reducing the average size of the gene clusters (3.73 ± 3.75 vs 3.44 ± 3.53; Figure S8CDE). In addition the variation of the COG scores within the gene clusters decreased, suggesting that fewer housekeeping genes would be added to detected biosynthetic gene clusters (Figure S8F).

Overlapping gene clusters were fused, resulting in 7.18 *10^5^ gene clusters. To organize the results, all clusters were paired if their protein domain content was similar (Jaccard index of protein domains; cutoff: 0.5) and at least one of their predicted precursors showed sequence similarity (NCBI blastp; bitscore cutoff: 30). These cutoffs were used to distinguish between different RiPP subclasses (Figure S9). Clustering these pairs with MCL created 45,727 ‘families’ of gene clusters, containing 312,163 gene clusters, while the remaining 406,105 gene clusters were left ungrouped.

Analysis of overlap between decRiPPter clusters and BGCs predicted by antiSMASH revealed that 5,908 clusters overlapped, constituting 78% of antiSMASH hits, but only 0.8% of decRiPPter clusters (Table 1, row 2). To further narrow down our results, we applied several filters to increase the saturation of RiPP BGCs in our dataset. A mild filter, limiting the average COG score to 0.25 and requiring two biosynthetic genes and a gene encoding a transporter, increased the fraction of overlapping RiPP BGCs to 7.8% (Table 1, row 3). When only clusters associated with genes for a predicted peptidase and a predicted regulator were considered, and the average COG score was limited to 0.1, the fraction increased further to 14.4% (Table 1, row 4). While many antiSMASH RiPP BGCs were filtered out in the process (and, by extension, many unknown RiPP BGCs were likely also filtered out this way), we felt our odds of discovering novel RiPP families were highest when focusing on the dataset with the highest fraction of RiPP BGCs, and therefore applied the strict filter. The remaining 2,471 clusters of genes were clustered as described above. Since our efforts were aimed at finding new gene cluster families, we discarded groups of clusters with fewer than three members, leaving 1,036 gene clusters in 187 families. Families in which more than half of the gene clusters overlapped with antiSMASH non-RiPP BGCs were discarded as well, leaving only known RiPP families and new candidate RiPP families (893 gene clusters, 151 families; Figure 2).

**Figure 2.**
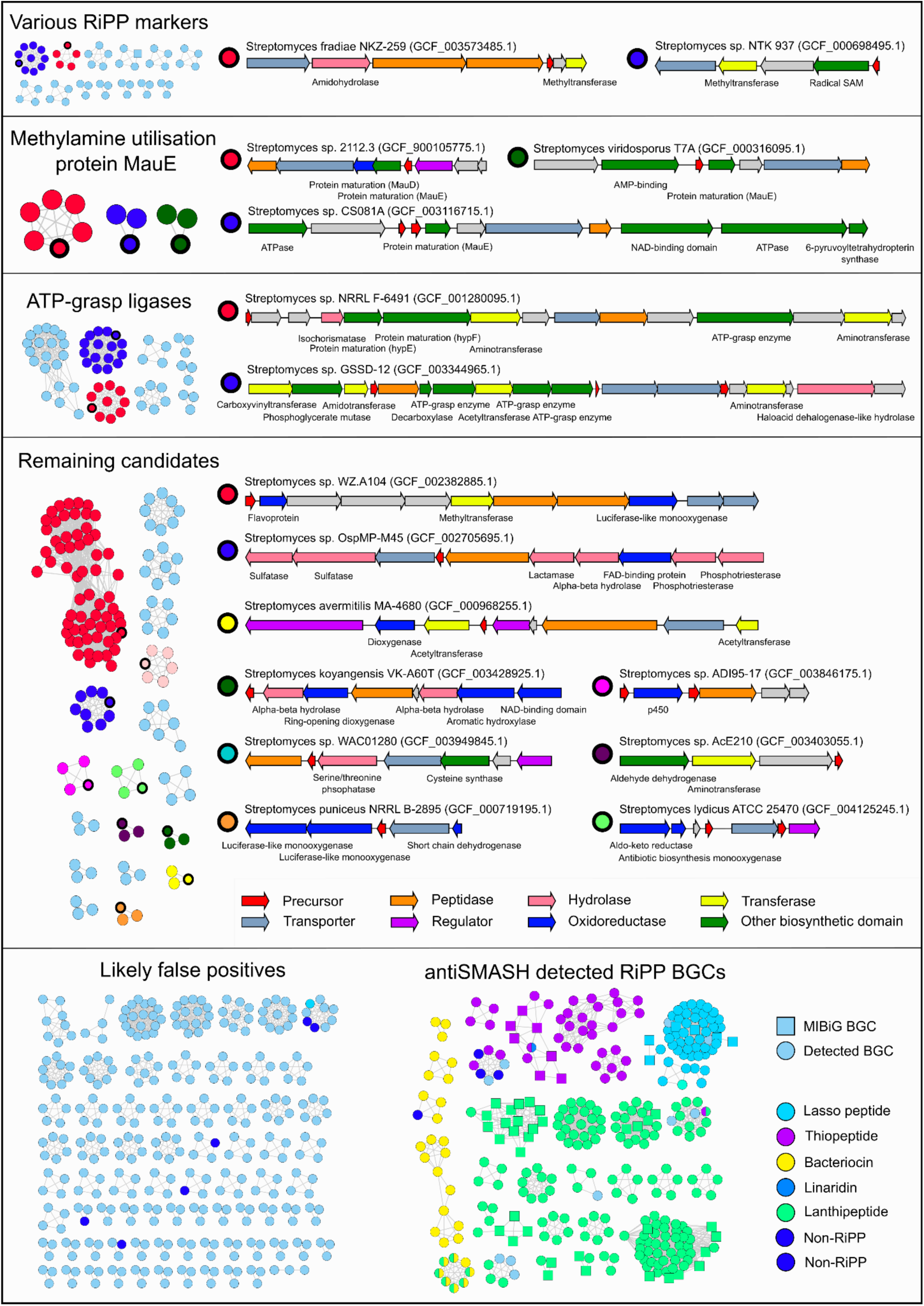
decRiPPter finds 42 candidate RiPP families with a large variety of encoded modifying enzymes and precursors. Gene clusters found in 1,295 *Streptomyces* genomes were passed through a strict filter and grouped together (see main text). Arrow colors indicate enzyme family of the product, and the description of gene products is given below the arrows. Roughly a third of the remaining candidates overlapped with or were similar to RiPP BGCs predicted by antiSMASH. Another third of the remaining candidates were discarded as likely false positives (see main text). Of the remaining 42 candidate RiPP families, 15 example gene clusters are displayed.

Roughly a third (272) of the remaining gene clusters were members of known families of RiPPs, including lasso peptides, lanthipeptides, thiopeptides, bacteriocins and microcins. In addition, many of the other candidate clusters (55) contained genes common to known RiPP BGCs, such as those encoding YcaO cyclodehydratases and radical SAM-utilizing proteins (Figure 2). These gene clusters were not annotated as RiPP gene clusters by antiSMASH, but the presence of these genes alone or in combination with a suitable precursor can be used as a lead to find novel RiPP gene clusters^24,29^.

Each remaining family of gene clusters was manually investigated to filter out likely false positives from the candidates. Common reasons to discard gene clusters were functional annotations of candidate precursors as having a non-precursor function (e.g. homologous to ferredoxin or LysW^55^), annotations of the genes within a gene cluster related to primary metabolism (e.g. genes for cell-wall modifying enzymes), or other abnormalities (e.g. large intergenic gaps or very large gene cluster of more than 40 genes). Several modifying enzymes belonging to the candidate families were homologous to gene products involved in primary metabolism, such as 6-pyruvoyltetrahydropterin synthase or phosphoglycerate mutase. Given the low distribution (COG scores) of the genes encoding these enzymes, it seemed more likely to us that they were adapted from primary metabolism to play a role in secondary metabolism^17^. We therefore only discarded a gene cluster family if multiple clear relations to a known pathway were found. The remaining 42 candidate families were further grouped together into broader classes depending on whether a common enzyme was found (Figure 2).

A large group of families all contained one or more genes for ATP-grasp enzymes. ATP-grasp enzymes are all characterized by a typical ATP-grasp-fold, which binds ATP, which is hydrolyzed to catalyze a number of different reactions. As such, these enzymes have a wide variety of functions in both primary and secondary metabolism, and their genes are present in a many different genomic contexts^56^. Involvement of ATP-grasp enzymes in RiPP biosynthesis has been reported for both microviridin^57^ and pheganomycin^23^, where they catalyze macrocyclization and peptide ligation, respectively. The ATP-grasp enzymes involved in the biosynthesis of these products did not show direct similarity to any of the ATP-grasp ligases of these candidates, however, suggesting that these belong to yet to be uncovered biosynthetic pathways.

Among the candidate families were three families that contained homologs to *mauE*, and one that additionally contained a homolog of *mauD*. The proteins encoded by these genes, along with other proteins encoded in the *mau* gene cluster, are known to be involved in the maturation of of methylamine dehydrogenase, which is required for methylamine metabolism. MauE in particular has been speculated to play a role in the formation of disulfide bridges in the β-subunit of the protein, while the exact function of MauD remains unclear^58^. As no other orthologs of the *mau* cluster were found within the genomes of *Streptomyces sp.* 2112.3, *Streptomyces viridosporus* T7A or *Streptomyces sp.* CS081A, it is unlikely that these proteins carry out this function. Rather, the presence of these genes in a putative RiPP BGC suggests that they play a role in modification of RTEs or RiPP precursors. Supporting this hypothesis, each of these gene clusters contained a gene predicted to a encode for a precursor containing at least eight cysteine residues (Table S3).

Similarly, homologs of *hypE* and *hypF* were detected in a gene cluster containing another gene encoding an ATP-grasp ligase. Genes encoding these proteins are typically part of the *hyp* operon, which is involved in the maturation of hydrogenase. Specifically, the two proteins cooperate to synthesize a thiocyanate ligand, which is transferred onto an iron center and used as a catalyst^59^. No other homologs of genes in the *hyp* operon were detected, however, suggesting that these protein-coding genes have adopted a novel function.

The remaining 18 families could not be grouped under a single denominator, nor could any single enzyme be found that clearly distinguished these groups as RiPP or non-RiPP BGCs. A wide variety of enzymes was found to be encoded by these gene clusters, including p450 oxidoreductases, flavoproteins, aminotransferases, methyltransferases and phosphatases. In addition (and in line with features dominant in the positive training set), the predicted precursor peptides were often rich in cysteine, serine and threonine residues (Table S3), which contain reactive hydroxyl and sulfide moieties and are present in precursors of various known RiPP subclasses.

All candidate gene clusters presented here carry the features we selected, typical of RiPP BGCs: a low frequency of occurrence among the scanned genomes, a suitable precursor peptide, candidate modifying enzymes, transporters, regulators and peptidases. However, many known RiPP BGCs were removed, suggesting that there may be more uncharacterized RiPP families among the gene clusters we discarded. While the complete dataset could not be covered here, the command-line application of decRiPPter has been set up to allow users to set their own filters. In addition, decRiPPter runs are visualized in an HTML output, in which the results can be further browsed and filtered by Pfam domains and other criteria, allowing users to find candidate families according to their preferences. The results from this analysis of the strict and the mild filter is available at http://www.bioinformatics.nl/~medem005/decRiPPter_strict/index.html and http://www.bioinformatics.nl/~medem005/decRiPPter_mild/index.html, respectively.

### Discovery of a novel family of lanthipeptides

To validate the capacity of decRiPPter to find novel RiPP subclasses, we set out to experimentally characterize one of the candidate families (Figure 2; Other; red marker). Gene clusters belonging to this family shared several genes encoding flavoproteins, methyltransferases, oxidoreductases and occasionally a phosphotransferase. Importantly, the predicted precursor peptides encoded by these putative BGCs showed clear conservation of the N-terminal region, while varying more in the C-terminal region (Figure S10). This distinction is typical of RiPP precursors, as the N-terminal leader peptide is used as a recognition site for modifying enzymes, while the C-terminal core peptide can be more variable^20^.

One of the gene clusters belonging to this candidate family was identified in *Streptomyces pristinaespiralis* ATCC 25468 (fig 3A; Table 2). *S. pristinaespiralis* is known for the production of pristinamycin, and was selected for experimental work since the strain is genetically tractable^60,61^. The gene cluster was named after its origin (*spr*: *Streptomyces pristinaespiralis* RiPP), and the genes were named after their putative function.

**Table 2.**
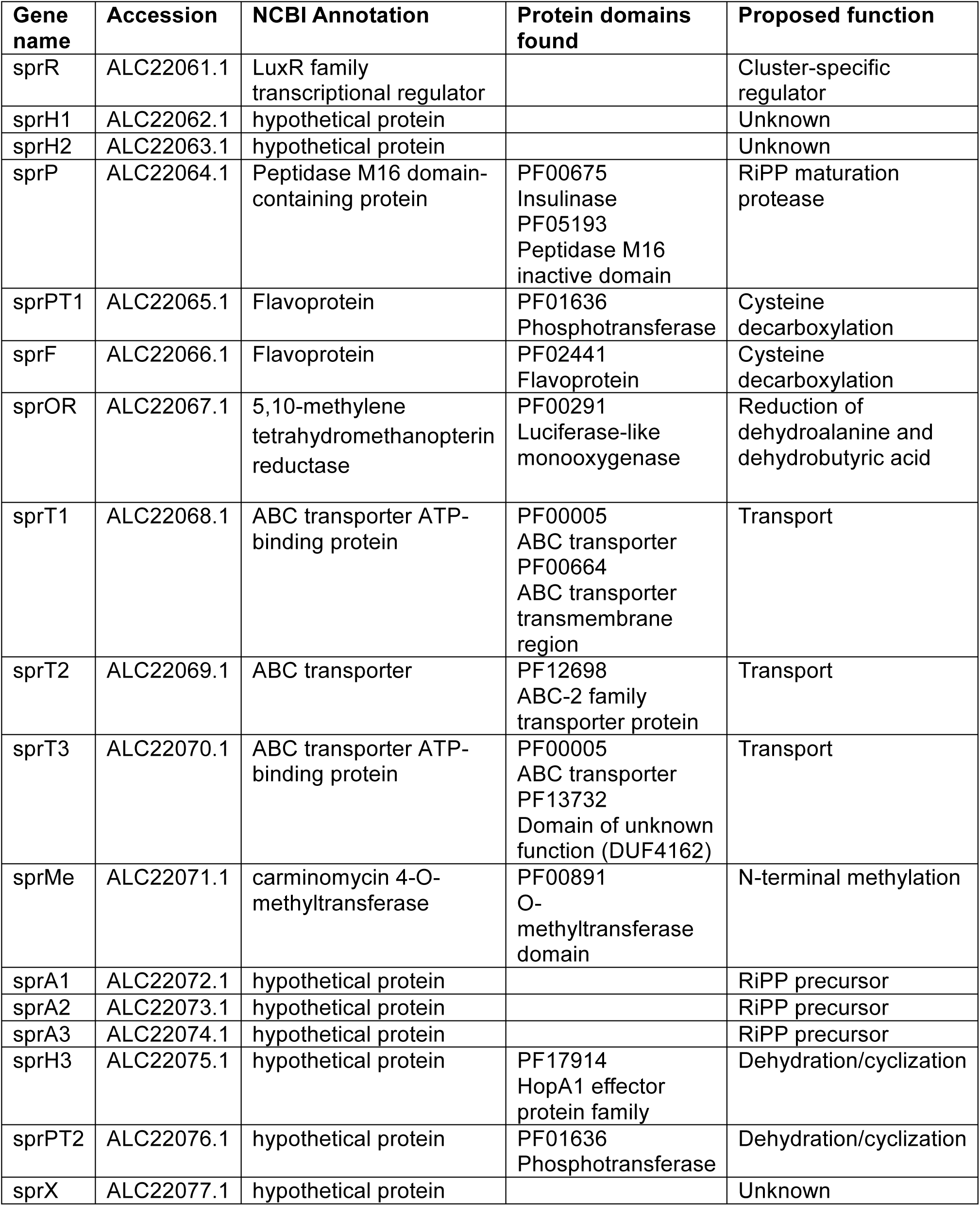
Annotation of the *Streptomyces pristinaespiralis* RiPP (*spr*) gene cluster.

| Gene name | Accession | NCBI Annotation | Protein domains found | Proposed function |
| --- | --- | --- | --- | --- |
| sprR | ALC22061.1 | LuxR family transcriptional regulator |  | Cluster-specific regulator |
| sprH1 | ALC22062.1 | hypothetical protein |  | Unknown |
| sprH2 | ALC22063.1 | hypothetical protein |  | Unknown |
| sprP | ALC22064.1 | Peptidase M16 domain-containing protein | PF00675<br>Insulinase<br>PF05193<br>Peptidase M16 inactive domain | RiPP maturation protease |
| sprPT1 | ALC22065.1 | Flavoprotein | PF01636<br>Phosphotransferase | Cysteine decarboxylation |
| sprF | ALC22066.1 | Flavoprotein | PF02441<br>Flavoprotein | Cysteine decarboxylation |
| sprOR | ALC22067.1 | 5,10-methylene tetrahydromethanopterin reductase | PF00291<br>Luciferase-like monooxygenase | Reduction of dehydroalanine and dehydrobutyric acid |
| sprT1 | ALC22068.1 | ABC transporter ATP-binding protein | PF00005<br>ABC transporter<br>PF00664<br>ABC transporter transmembrane region | Transport |
| sprT2 | ALC22069.1 | ABC transporter | PF12698<br>ABC-2 family transporter protein | Transport |
| sprT3 | ALC22070.1 | ABC transporter ATP-binding protein | PF00005<br>ABC transporter<br>PF13732<br>Domain of unknown function (DUF4162) | Transport |
| sprMe | ALC22071.1 | carminomycin 4-O-methyltransferase | PF00891<br>O-methyltransferase domain | N-terminal methylation |
| sprA1 | ALC22072.1 | hypothetical protein |  | RiPP precursor |
| sprA2 | ALC22073.1 | hypothetical protein |  | RiPP precursor |
| sprA3 | ALC22074.1 | hypothetical protein |  | RiPP precursor |
| sprH3 | ALC22075.1 | hypothetical protein | PF17914<br>HopA1 effector protein family | Dehydration/cyclization |
| sprPT2 | ALC22076.1 | hypothetical protein | PF01636<br>Phosphotransferase | Dehydration/cyclization |
| sprX | ALC22077.1 | hypothetical protein |  | Unknown |

**Table 3.** Co-occurrence of genes found in the *spr* gene cluster with homologs of *sprPT* in the analyzed 1,295 *Streptomyces* strains.

| Gene name | Co-occurrence with *sprPT*<br>(percentage) |
| --- | --- |
| *sprH3* | 99.49 |
| *sprMe* | 20 |
| *sprT1* | 35.38 |
| *sprT2* | 12.31 |
| *sprT3* | 12.82 |
| *sprOR* | 64.62 |
| *sprF1* | 39.5 |
| *sprF2* | 68.72 |
| *sprP* | 38.5 |
| *sprH1* | 9.0 |
| *sprH2* | 2.0 |
| *sprR* | 28.5 |
| *sprA1* | 1.03 |
| *sprA2* | 1.03 |
| *sprA3* | 16.92 |

**Figure 3.**
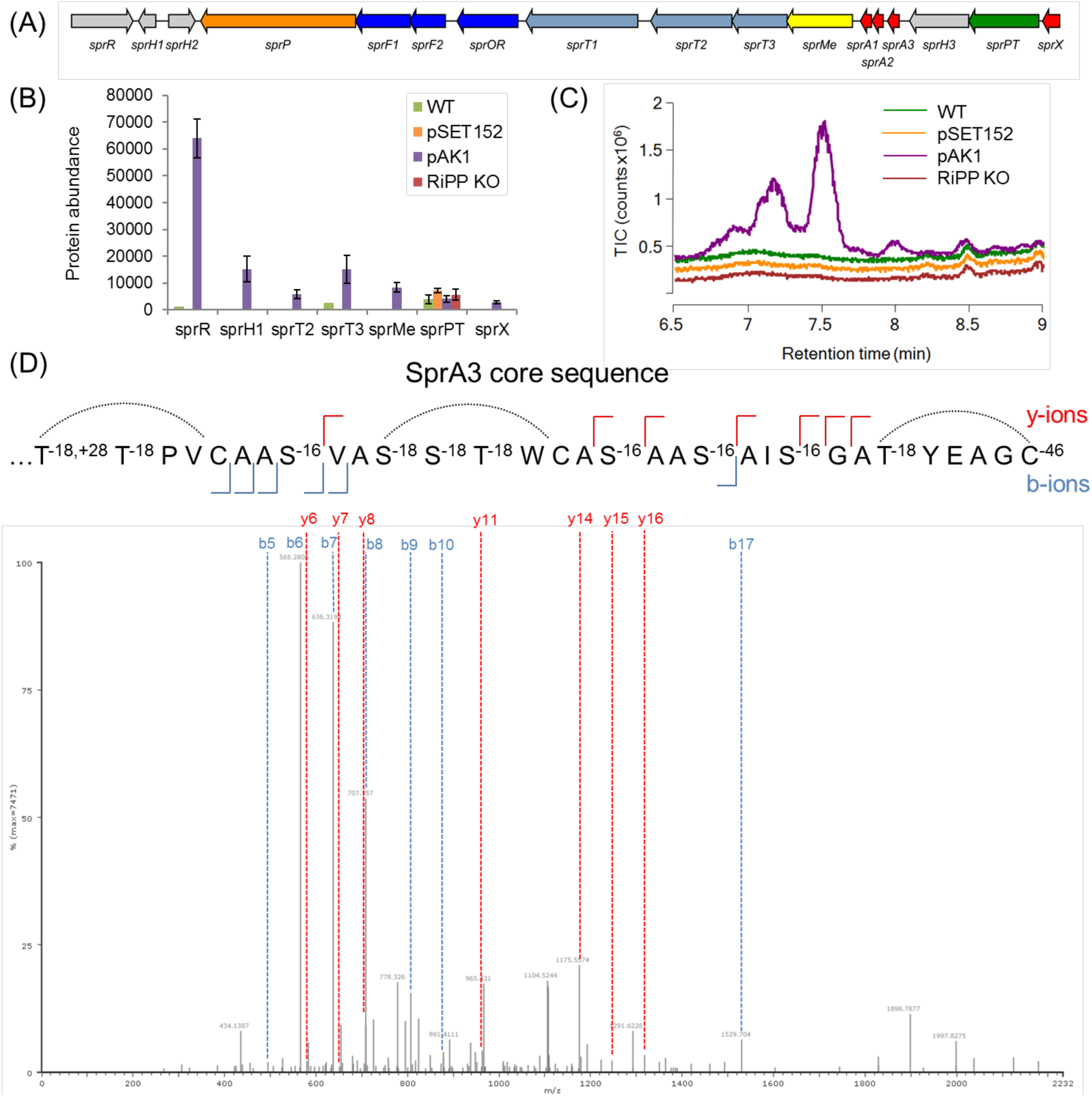
The *Streptomyces pristinaespiralis RiPP* (spr) gene cluster produces a highly modified RiPP. A) The *spr* gene cluster encodes three putative precursors, three transporters, a peptidase and an assortment of modifying enzymes (see Table 1). B) Protein abundance of the products of the *spr* gene cluster in *S. pristinaespiralis* ATCC 25468 and derived strains. Strains were grown in NMMP and samples were taken after 2 and 7 days. Enhanced expression of the regulator (from construct pAK1) resulted in the partial activation of the gene cluster. Genes that could not be detected are not illustrated. C) Chromatogram of crude extracts from strains grown under the same conditions as under A), samples after 7 days. Several peaks were detected in the extract from the strain with expression construct pAK1 between 7 and 8 minutes. C) b and y ions detected from one of the predominant peaks found in the crude extract (corresponding to monoisotopic mass of 2703.235 Da). The fragmentation pattern could be matched to the sprA3 precursor.

The gene cluster contains four genes encoding putative precursor peptides, although only three of the peptides (SprA1-A3) showed similarity to each other and to the other peptides in the same family (Figure S10). The fourth predicted precursor peptide (encoded by s*prX*) did not align with any of the other peptides and was assumed to be a false positive. The products encoded by *sprA1* and *sprA2* were highly similar to one another compared to the *sprA3* gene product. Occurrence of two distinct genes for precursors within a single RiPP BGC is typical for two-component lanthipeptides^62^.

Most of the modifying enzymes present in the gene cluster had not previously been implicated in RiPP biosynthesis. The predicted *sprF2* gene product, however, shows high similarity to cysteine decarboxylases such as EpiD and CypD. These enzymes decarboxylate C-terminal cysteine residues, which is the first step in the formation of C-terminal loop structures called S-[(Z)-2-aminovinyl]-D-cysteine (AviCys) and S-[(Z)-2-aminovinyl]-(3S)-3-methyl-D-cysteine (AviMeCys)^63^. Several RiPP classes have been reported with this modification, including lanthipeptides, cypemycins and thioviridamides, although they are only consistently present in cypemycins and thioviridamides. This type of modification is less common among lanthipeptides, with only nine out of 120 lanthipeptide gene clusters in MIBiG encoding the required decarboxylase. Cysteine-decarboxylating genes are also present in non-RiPP gene clusters (Table S4) and are also associated with other metabolic pathways^64^.

A more detailed comparison with the gene clusters in MIBiG showed that two more genes from the thioviridamide gene cluster were homologous to two genes encoding a predicted phosphotransferase (*sprPT*) and a hypothetical protein (*sprH3*), respectively. Taken together with the homologous cysteine decarboxylase, it appeared that our gene cluster was distantly related to the thioviridamide gene cluster^65^. Thioviridamide-like compounds are primarily known for thioamide residues, for which a TfuA-associated YcaO is thought to be responsible^29,66^. However, a YcaO homologue was not encoded by the gene cluster, making it unlikely that this gene cluster should produce thioamide-containing RiPPs.

Two strains were created to help determine the natural product specified by the BGC. For the first strain, the entire gene cluster was replaced by an apramycin resistance cassette (aac3(IV)) by homologous recombination with the pWHM3 vector^67^ (*spr*::apra). In case the gene cluster was natively expressed, this strain should allow for easy identification of the natural product by comparative metabolomics. In the second approach, we sought to activate the BGC in case it was not natively expressed. To this end, we targeted the cluster-situated *luxR*-family transcriptional regulatory gene *sprR*. The *sprR* gene was expressed from the strong and constitutive *gapdh* promoter from *S. coelicolor* (p_gapdh_) on the integrative vector pSET152^68^. The resulting construct (pAK1) was transformed to *S. pristinaespiralis* by protoplast transformation.

To assess the expression of the gene cluster in the transformants, we analyzed changes in the global expression profiles in 2 days and 7 days old samples of NMMP-grown cultures using quantitative proteomics (Figure 3B). Aside from the regulator itself, six out of the sixteen other proteins were detected in the strain containing expression construct pAK1, while only SprPT could be detected in the strain carrying the empty vector pSET152. SprPT was also detected in the proteome of *spr*::apra, however, indicating a false positive. In the wild-type strain, SprT3 and SprR were detected, but only in a single replicate and at a much lower level. Overall, these results suggest that under the chosen growth conditions the gene cluster was expressed at very low amounts in wild-type cells, and was activated when the expression of the likely pathway-specific regulatory gene was enhanced. This makes *spr* a likely cryptic BGC.

To see if a RiPP was produced, the same cultures used for proteomics were separated into mycelial biomass and supernatant. The biomass was extracted with methanol, while HP20 beads were added to the supernatants to absorb secreted natural products. Analysis of the crude methanol extracts and the HP20 eluents with HPLC-MS revealed several peaks eluting between 5.5 and 7 minutes in the methanol extracts (fig 3C), which were not found in extracts from wild-type strain or the strain containing the empty vector. Feature detection with MZMine followed by statistical analysis with MetaboAnalyst revealed seven unique peaks, with m/z between 707.3534 and 918.0807 (Figure S11). The isotope patterns of these peaks showed that the six of the corresponding compounds were triply charged. Careful analysis of derivative peaks with mass increases consistent with Na- or K-addition, led to the conclusion that these peaks corresponded to the [M+3H]^3+^ adduct, suggesting a monoisotopic masses in the range of 2,604.273 and 2,754.242 Da. The highest signal came from the compound with monoisotopic mass of 2,703.245. Four of the other masses seemed to be related to this mass, as they were different in increments of 4, 14, or 16 Da (Table S5). We therefore reasoned that the mass of 2,703.245 Da was the final product, while others were incompletely processed peptides.

To further verify that the identified masses indeed belonged to the RiPP precursors in our gene cluster, we first removed the apramycin resistance cassette from Spr::apra using the pUWLCRE vector^69^, creating strain Δ*spr*. The expression construct pAK1 and an empty pSET152 vector were transformed to the strain Δ*spr*. When these strains were grown under the same conditions, the aforementioned peaks were not detected, further suggesting that indeed they belonged to products of this gene cluster (Figure S12).

Most masses were detected in only low amounts. In order to resolve this, we created a similar construct as pAK1, but this time using the low-copy shuttle vector pHJL401 as the vector^70^. The plasmid pAK2 was introduced into *S. pristinaespiralis* and the transformants grown in NMMP for 7 days. Extraction of the mycelial biomass with methanol resulted in a higher abundance of the masses previously detected (Figure S13). Consistent with the MS profiles of pAK1 transformants, also pAK2 transformants produced an abundant peak corresponding to a monoisotopic mass of 2,703.245 Da, as well as a second peak corresponding to a monoisotopic mass of 2,553.260 Da. Most of the other masses could be related to one of these two masses, suggesting these are the final products, related to two distinct precursors (Tables S5 and S6).

We then performed MS-MS analysis of the extracts of the pAK2 transformants to identify the metabolites and their expected modifications, such as Avi(Me)Cys moieties. The fragmentation pattern of the mass of 2,703.245 Da could be assigned to the sprA3 precursor, when several modifications were applied (Figure 3D, Table S7). Similarly, fragments with a mass of 2,553.260 could be matched to the SprA2 precursors considering the same modifications (Figure S14; Table S8).

Among the predicted modifications were N-terminal methylation, which was supported by the presence of the methyltransferase *sprMe* in the gene cluster. Secondly, the C-terminal cysteine was predicted to have undergone oxidative decarboxylation (−46 Da), as expected based on the presence of the gene *sprF2* in the gene cluster. In addition, many of the serines and threonines could only be matched when their masses were altered by −16 or −18 Da. These mass differences are typical of dehydration (−18 Da) of the residues to dehydroalanine and dehydrobutyric acid. Reduction of these dehydrated amino acids (+2 Da) would then give rise to alanine and butyric acid residues, a modification which has been reported for lanthipeptides^71^.

To test for the presence of dehydrated serines and threonines, we treated the purified product with dithiothreitol (DTT), which covalently attaches to these residues via 1,4 nucleophilic addition^72^. Treatment with DTT resulted in the addition of up to two adducts, showing the presence of dehydrated residues, although one fewer than expected (Figure S15). The fact that two of the dehydrated residues are adjacent to one another may have resulted in steric hindrance, preventing full conversion.

Surprisingly, no fragments were found of the residues S^-18^S^-18^T^-18^WC in the center of SprA3, or for the N-terminal T^-18, +28^T^-18^PVC region. Considering the other modifications typical of lanthipeptides, we hypothesized the presence of thioether crosslinks between the dehydrobutyric acids and cysteines. To find further support for this hypothesis, we treated the purified product of SprA3 with iodoacetamide (IAA). Iodoacetamide alkylates free cysteines, while cysteines in thioether bridges remain unmodified^73^. In agreement with our hypothesis, treatment with iodoacetamide did not affect the observed masses, despite the presence of three cysteines in the peptide (Figure S10). In addition, we hydrolyzed the purified peptide with 6M HCl at 110°C for 24h. Under these conditions, the amide bond should be hydrolyzed, while the thioether bond should be unaffected^74^. The resulting mixture of amino acids both contained masses corresponding to a cysteine linked to either a dehydrated serine, or to a twice methylated, dehydrated threonine (Table S10). The C-terminal predicted AviMeCys was not detected, although this may be explained by the presence of the alkene in the moiety, which are likely to react under acidic conditions.

Many of the other masses found were higher when compared to the product of SprA3 by increments of 16 Da, suggesting that the peptide was incompletely processed. The fragmentation patterns of these masses could not be unambiguously resolved (Figure S16). An unmodified serine or threonine could occur at several places within the precursor, and each of the possible outcomes would likely give rise to compounds with identical mass and very similar hydrophobic properties, which would not be separated properly. Overall, these results further reinforce the idea that the compound with monoisotopic mass of 2,703.245 Da belongs to the fully modified product, while the others are derived from it.

### The *sprH3/sprPT* gene pair is present in a wide variety of RiPP-like contexts

Taken together, we have shown that the SprA3 precursor contained a number of posttranslational modifications that are typical of lantibiotics. The conversion of serine/threonine to alanine/butyric acid via reduction, the creation of an AviCys moiety and the crosslinks to form thioether bridges are all found in lanthipeptides, and are dependent on dehydration of serine and threonine residues. Four different sets of enzymes, called LanBC, LanM, LanKC and LanKL can catalyze these reactions in the biosynthesis of lanthipeptides and are used to designate the lanthipeptide type.

As stated before, no members of any of these enzyme families were found to be encoded by the gene cluster studied. However, *sprH3* and *sprPT* showed homology to two uncharacterized genes of the thioviridamide BGC. Thioviridamide contains an AviCys moiety, the formation of which requires a dehydrated serine residue. The enzymes responsible for dehydration and subsequent cyclization have not been identified yet^65,75^. Since both gene clusters share a common modification for which the enzyme is unknown, we hypothesized that *sprH3* and *sprPT* should be responsible for dehydration and cyclization, and thus are hallmarks for a new lanthipeptide subtype, which we designate type V.

Lanthipeptide core modifying enzymes catalyze the most prominent reaction in lanthipeptide maturation, and as such, are present in many different genetic contexts^54^. To validate that SprH3 and SprPT are the sought-after modifying enzymes, we studied the distribution of the *SprH3*/*PT* gene pair across *Streptomyces* genomes analyzed by decRiPPter. Using CORASON^76^ with the s*prPT* gene as a query yielded 195 homologs in various gene clusters (Figure 4). The *sprPT/sprH3* gene pair was completely conserved across all gene clusters for which an uninterrupted contig of DNA was available., strongly supporting their functional interaction and joint involvement. Using the *sprH3* gene as a query yielded similar results (data not shown). A total of 391 orthologs of the gene pair were found outside *Streptomyces*, particularly in Actinobacteria (219) and Firmicutes (161; Figure S17). Distantly similar homologs of the gene pair were also identified in Cyanobacteria, Plantomycetes and Proteobacteria.

**Figure 4.**
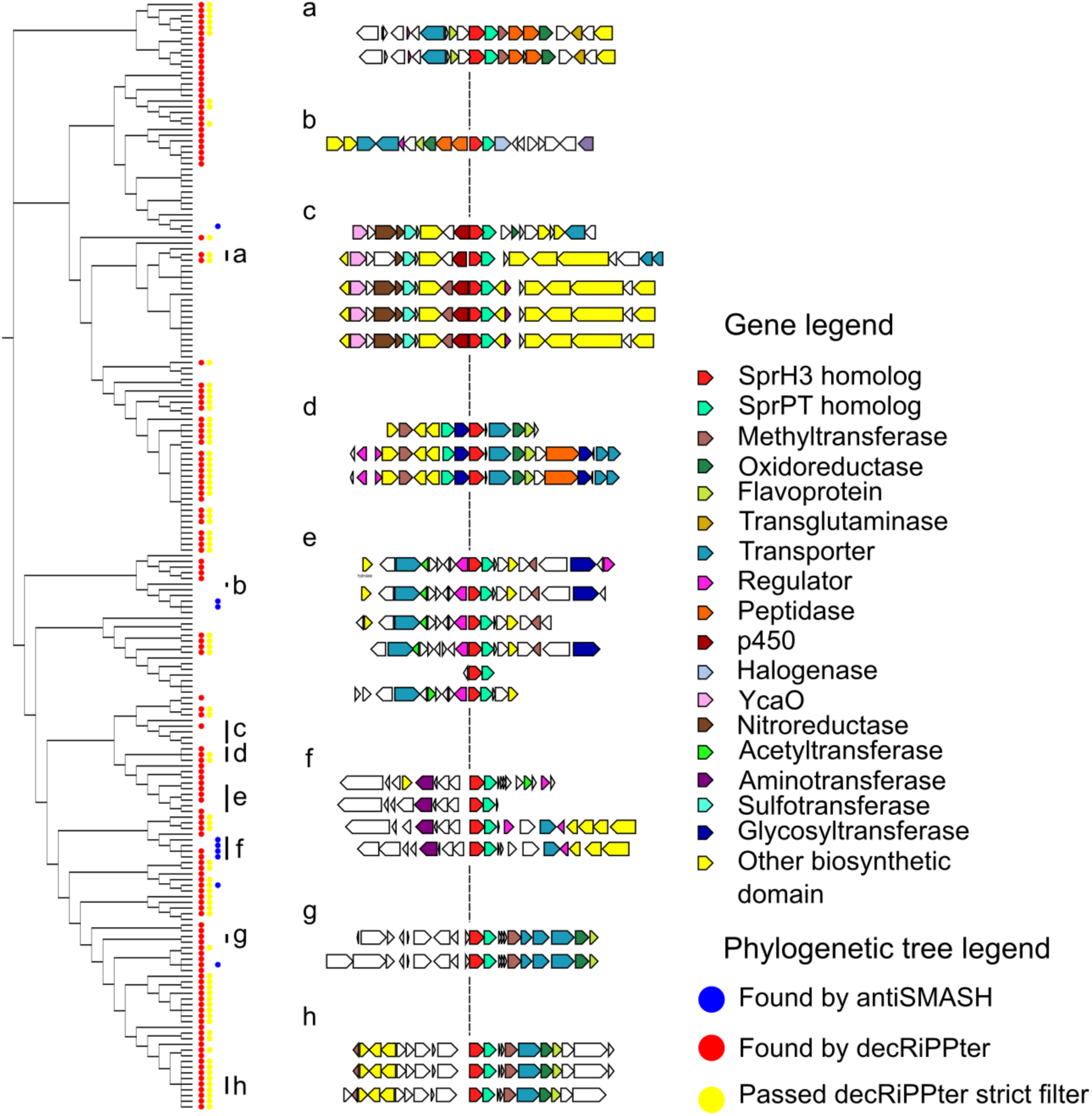
Orthologs of *sprPT* and *sprH3* cooccur in a wide variety of genetic contexts. (Left side) Phylogenetic tree of gene clusters containing homologs of *sprPT* and *sprH3*, visualized by CORASON^76^. A red dot indicates that the genes were present in a gene cluster found by decRiPPter, a yellow dot that it passed the strict filter (see Table 1 for details). A blue dot indicates overlap with a BGC identified by antiSMASH. (Right side) Several gene clusters with varying genetic contexts are displayed. Group (g) represents the query gene cluster. The genetic context varies, while the gene pair itself is conserved. Color indicates predicted enzymatic activity of the gene products as described in the legend.

Among the 195 identified gene clusters in *Streptomyces*, the majority (131) overlapped with a gene cluster detected by decRiPPter, indicating that the gene pair was within short intergenetic distance from predicted precursor gene in the same strand orientation. A large fraction (80) also passed the strictest filtering (see Table I), showing that among these gene clusters were many encoding biosynthetic machinery, peptidases and regulators. In contrast, only nine of the gene clusters overlapped with a BGC identified by antiSMASH. Four of these showed the gene pair in apparent operative linkage with a bacteriocin gene cluster, marked as such by the presence of a DUF692 domain, which is often associated with small prepeptides such as methanobactins. Another four gene clusters detected by decRiPPter were only overlapping due to the gene pair being on the edge of a neighboring gene cluster.

The genetic context of the gene pairs showed a wide variation (Figure 4, right side). While some gene clusters were mostly homologous to the *spr* gene cluster (Figure 4, group g-h), others shared only a few genes (groups a and d), and some only shared the gene pair itself (groups b, c and e). Many other predicted enzyme families were found to be encoded inside these gene clusters, including YcaO-like proteins, glycosyltransferases, sulfotransferases and aminotransferases. The large variation in genetic contexts combined with the clear association with a predicted precursor indicates that this gene pair likely plays a role in many different RiPP-associated genetic contexts, supporting their proposed role as a core gene pair.

Furthermore, we searched for genes encoding enzymes whose functions are dependent on a lanthipeptide dehydration in their substrate, to find if they were associated with the *sprPT/sprH3* gene pair. Both within and outside *Streptomyces*, homologs of *sprF1* and *sprF2* were often found associated with the gene pair (*sprF1*: 251/586; 40.1%; *sprF2*: 281/586; 48.0%; Table S11). Another modification dependent on the presence of dehydrated serine and threonine residues is the conversion of these to alanine and butyric acid, respectively, catalyzed by LtnJ and CrnJ^71^. Outside *Streptomyces*, the genomic surroundings of the *sprPT/sprH3* gene pair occasionally contained homologs of the *ltnJ* gene (40/391; 10.1%), further implying that these genes carry out the canonical dehydration reactions.

A similar modification was observed for SprA2 and SprA3, despite that no homologs of the genes encoding LtnJ or CrnJ were identified within the *spr* gene cluster. However, *sprOR* encodes a putative oxidoreductase, and thus candidates for this modification. Supporting this, orthologs of *sprOR* were found frequently associated with either canonical lanthipeptide BGCs or the *sprPT/sprH3* gene pair (lanthipeptide: 124/462; *sprPT/sprH3*: 137/462; Table S10). One of these lanthipeptide BGCs showed high homology to the lacticin 3147 BGCs from *Lactococcus lactis*. Lacticin 3147 contains several D-alanine residues as a result of conversion of dehydrated serine residues^77^. While all the genes, including the precursors, were well conserved between the two gene clusters, the *ltnJ* gene had been replaced by an *sprOR* homolog, suggesting that their gene products catalyze similar functions (Figure S18).

### Conclusion and final perspectives

The continued expansion of available genomic sequence data has allowed for discovery of large reservoirs of natural product BGCs, fueled by sophisticated genome mining methods. These methods must make tradeoffs between novelty and accuracy^12^. Tools primarily aimed at accuracy reliably discover large numbers of known natural product BGCs, but are limited by specific genetical markers. On the other hand, while tools aimed at novelty may discovery new natural products, these tools have to sacrifice on accuracy, resulting in a larger amount of false positives.

Here, we take a new approach to natural product genome mining, aimed specifically at the discovery of novel types of RiPPs. To this end, we built decRiPPter, an integrative approach to RiPP genome mining, based on general features of RiPP BGCs rather than selective presence of specific types of enzymes and domains. To increase the accuracy of our methods, we base detection of the RiPP BGCs on the one thing all RiPP BGCs have in common: a gene encoding a precursor peptide. With this method, we identify 42 candidate novel RiPP families, mined from only 1,295 *Streptomyces* genomes. These families are undetected by antiSMASH, and show no clear markers identifying them as belonging to previously known RiPP BGC classes. While the approach to RiPP genome mining taken here inevitably gives rise to a higher number of false positives, we feel that such a ‘low-confidence / high novelty’ approach^12^ is necessary for the discovery of completely novel RiPP families. Additionally, users are able to set their own filters for the identified gene clusters, allowing them to search candidate RiPP families containing specific enzymes or enzyme types within a much more confined search space compared to manual genome browsing.

The product of one of the candidate classes was characterized as the first member of a new class of lanthipeptides (termed ‘type V’) that was not detected by any other RiPP genome mining tool. Variants of this gene cluster are widespread across *Streptomyces* species, further expanding one of the most widely studied RiPP families. In addition, two proposed core genes were used to expand the family by finding additional homologs in *Actinobacteria* and *Firmicutes*. Taken together, this work shows that known RiPP families only cover part of the complete genomic landscape, and that many more RiPP families likely remain to be discovered, especially when expanding the search space to the broader bacterial tree of life.

