## Supplemental Document 1 for "Integration of machine learning and pan-genomics expands the biosynthetic landscape of RiPP natural products"

### Index

Material and methods ........ 2
 Experimental ........ 2
 Bioinformatics ........ 5
Figure S1. Gene cluster formation pipeline ........ 10
Figure S2. COG score calculation pipeline ........ 11
Figure S3. Effects of trueCOGs on cutoffs and genome sizes ........ 12
Figure S4. COG score distribution in *Streptomyces* ........ 13
Figure S5. COG scores of BUSCO genes ........ 14
Figure S6. COG scores of antiSMASH BGCs ........ 15
Figure S7. Distribution of SVM scores in *Streptomyces* ........ 16
Figure S8. Gene cluster formation in *Streptomyces* ........ 17
Figure S9. RiPP subclass clustering using precursor and domain similarity ........ 18
Figure S10. Alignment of precursors from the novel lanthipeptide RiPP family ........ 19
Figure S11. Heatmap comparing masses between pAK1-containing strains with WT ........ 20
Figure S12. LCMS chromatogram of Δspr containing pAK1 ........ 21
Figure S13. Comparison of extracts from strains containing pAK2 with WT ........ 22
Figure S14. Fragmentation pattern of mass 2553.260 Da matched to SprA2 ........ 23
Figure S15. Labeling of SprA3 with DTT and IAA ........ 24
Figure S16. Fragmentation pattern of mass 2719.245 Da matched to SprA3 derivative ........ 25
Figure S17. Comparison of lacticin-3147 BGC with *sprOR*-containing BGC ........ 26
Figure S18. Distribution of *sprPT/H3* gene pair outside *Streptomyces* ........ 27
Table S1. Plasmids used in this study ........ 28
Table S2. Primers used in this study ........ 29
Table S3. Precursor sequences of candidate RiPP families ........ 30
Table S4. Spread of the flavoprotein domain (PF02441) in MIBiG BGCs ........ 31
Table S5. Detected masses unique to pAK1-containing strains ........ 32
Table S6. Detected masses unique to pAK2-containing strains ........ 33
Table S7. Fragmentation ions of SprA3 ........ 34
Table S8. Fragmentation ions of SprA2 ........ 35
Table S9. Fragmentation ions of SprA3-derived compound ........ 36
Table S10. Detected amino acids from hydrolyzed SprA3 ........ 38
Table S11. Cooccurrence of dehydration-dependent genes with the *sprPT/sprH3* gene pair ........ 39
References ........ 40

#### Materials and methods

Experimental

*Bacterial strain and growth conditions*

*Streptomyces pristinaespiralis* ATCC 25468 was purchased from DSMZ (DSM number 40338). Media components were purchased from Thermo Fisher Scientific, Sigm-Aldrich or Duchefa Biochemie. For strain cultivation on solid media, *Streptomyces* spores were spread on mannitol soya flour agar (SFM; 20 g/L Agar, 20 g/L mannitol, 20 g/L soya flour, supplemented with tap water) prepared as described previously[71], and incubated at 30°C. Spores were harvested after 4-7 days of growth when the strain started to produce a grey pigment, by adding water directly to the plate and releasing the spores with a cotton swab. Spores were centrifuged and stored in 20% glycerol.

For cultivation in liquid media, 20-50 µL of a dense spore stock was inoculated into 100 mL shake flasks with coiled coils containing 20 mL of the medium of interest. For extractions, NMMP was used (0.60 mg/L $MgSO_4$, 5 mg/L $NH_4SO_4$, 5 g/L Bacto casaminoacids, 1 mL trace elements (1 g/L $ZnSO_4.7H_2O$, 1 g/L $FeSO_4.7H_2O$, 1 g/L $MnCl_2.4H_2O$, 1 g/L $CaCl_2$, anhydrous)), while for genomic DNA isolation, a 1:1 mixture of TSBS: YEME with 0.5% glycine and 5 mM $MgCl_2$ was used (TSBS: 30 g/L Bacto Tryptic Soy Broth, 100 g/L sucrose; YEME: Bacto Yeast Extract: 3 g/L, Bacto Peptone 5 g/L, Bacto Malt Extract 3 g/L, glucose 10 g/L, sucrose 340 g/L).

*E. coli* strains JM109 and ET8 were used for general cloning purposes and demethylation, respectively. Strains were cultivated in liquid LB and on LB-agar plates at 37°C.

*Molecular biology*

All materials and primers were purchased from Sigma-Aldrich or Thermo Fisher Scientific unless stated otherwise. Restriction enzymes and T4 ligase were purchased from NEB. Restriction and ligation protocols were followed as per manufacturer's description. For amplification of DNA fragments with PCR, Pfu polymerase was used. Primers were designed with $T_m$ of the annealing region roughly equal to 60°C. Standard PCR protocols consisted of 30 cycles (45 second DNA melting @ 95 °C, 45 second primer annealing @55°C-65°C, 60s-180s primer elongation @ 72°C), but PCR protocols were optimized where necessary.

*S. pristinaespiralis Spr::Ap* deletion mutants were created by replacing the gene cluster with an *aac(3)IV* apramycin resistance cassette via homologous recombination. The -1507/-39 and +135/+1641 regions upstream and downstream of the cluster were amplified by PCR with the spr_LF_F/spr_LF_R and spr_RF_F/spr_RF_R primer pairs (table SII), respectively, and inserted into the pWHM3-oriT vector (Table SI) into the EcoRI/HindIII sites. The *aac(3)IV* apramycin resistance cassette was inserted into the XbaI site, creating pAK3. pAK3 was transformed to *E.coli ET8* for DNA demethylation, purified, and transformed to *S. pristinaespiralis* by protoplast transformation. Transformation mixtures were plated out on R5, prepared as described earlier[71]. After 14-18 hours, the plates were overlaid with 1.2 mL $H_2O$ containing 10 µg thiostrepton and 25 µg apramycin. Three colonies were picked after 4 days of growth and spread onto SFM plates without added antibiotic to allow for homologous recombination. Colonies containing the correct phenotype (apramycin-resistant, thiostrepton-sensitive) were picked and the homologous recombination was confirmed by PCR, using the spr_del_check_F/spr_del_check_R primer pair.

Constructs for the overexpression of the *sprR* regulator were constructed as follows: the *sprR* gene was amplified from the genomic DNA of *S. pristinaespiralis* using the sprR_F/sprR_R primer pair, and placed into the EcoRI/XbaI site of the pSET152 vector. The -0/-457 upstream region of glyceraldehyde

3-phosphate dehydrogenase amplified from the genome of *S. coelicolor*, was obtained from previous studies[72,73] and inserted into the EcoRI site and the engineered NdeI site, placing it directly upstream of the *sprR* gene. To create vector pAK2, the entire region between the EcoRI and XbaI sites was excised and inserted into the pHJL401 vector.

*Extractions*

Strains were cultured in 100 mL shake flasks containing 20 mL NMMP, with coiled coils at 30°C for 7 days. 20 µg/mL thiostrepton was added to cultivate strains containing pHJL401. Mycelium was collected by centrifugation, washed twice with sterile MiliQ water and extracted with 5 mL methanol by shaking overnight at 4°C. The methanol was collected and centrifuged at 4°C to clear it of cellular debris and precipitates. The crude extracts were dried and weighed, and dissolved in methanol at a concentration of 1 mg/mL for further analysis.

*Peptide purification*

For large-scale extraction, the strain proved incapable of producing the desired compound when grown in large shake flasks. Therefore, 2L NMMP prepared as above was inoculated with 2.5 mL of a dense spore stock *S. pristinaespiralis* with pAK3, and split over one hundred 100 mL shake flasks. Thiostrepton was added as described above. The cultures were grown for 14 days, pooled together and extracted with an equivalent volume of butanol. The butanol extracted was then evaporated *in vacuo* to yield 1.7g of crude extract.

The resulting crude extract was adsorbed on silica gel 60 (40–60 µm, Sigma Aldrich), and dry loaded on a VLC column (3 × 30 cm) packed with the same material. The column was eluted with 200 mL fractions of a gradient comprised of (v/v): hexane, hexane–EtAc (1:1), EtAc, EtAc-MeOH (3:1), EtAc-MeOH (1:1), EtAc–MeOH (1:3), and finally MeOH. The fractions containing the compound of interest were pooled, concentrated and further purified using Waters preparative HPLC system comprised of 1525 pump, 2707 autosampler, and 2998 PDA detector. The pooled fraction (112.9 mg) was injected into a SunFire $C_{18}$ column (10 µm, 100 Å, 19 × 150 mm). The column was run at a flow rate of 12.0 mL/min, using solvent A (0.1% FA in $H_2O$) and solvent B (0.1% FA in ACN), and a gradient of 30–60% B over 20 min. HPLC purification was monitored at 254 nm, and eventually resulted in compound **1** (1.1 mg).

*LCMS analysis*

LC-MS/MS acquisition was performed using Shimadzu Nexera X2 UHPLC system, with attached PDA, coupled to Shimadzu 9030 QTOF mass spectrometer, equipped with a standard ESI source unit, in which a calibrant delivery system (CDS) is installed. The dry extracts were dissolved in MeOH to a final concentration of 1 mg/mL, and 2 µL were injected into a Waters Acquity HSS $C_{18}$ column (1.8 µm, 100 Å, 2.1 × 100 mm). The column was maintained at 30 °C, and run at a flow rate of 0.5 mL/min, using 0.1% formic acid in $H_2O$ as solvent A, and 0.1% formic acid in acetonitrile as solvent B. A gradient was employed for chromatographic separation starting at 5% B for 1 min, then 5 – 85% B for 9 min, 85 – 100% B for 1 min, and finally held at 100% B for 4 min. The column was re-equilibrated to 5% B for 3 min before the next run was started. The LC flow was switched to the waste the first 0.5 min, then to the MS for 13.5 min, then back to the waste to the end of the run. The PDA acquisition was performed in the range 200 – 600 nm, at 4.2 Hz, with 1.2 nm slit width. The flow cell was maintained at 40 °C.

The MS system was tuned using standard NaI solution (Shimadzu). The same solution was used to calibrate the system before starting. Additionally, a calibrant solution made from Agilent API-TOF reference mass solution kit was introduced through the CDS system, the first 0.5 min of each run, and the masses detected were used for post-run mass correction of the file, ensuring stable accurate mass measurements. System suitability was checked by including a standard sample made of 5 µg/mL paracetamol, reserpine, and sodium dodecyl sulfate; which was analyzed regularly in between the batch of samples.

All the samples were analyzed in positive (negative) polarity, using data-dependent acquisition mode. In this regard, full scan MS spectra (m/z 100 – 2000, scan rate 20 Hz) were followed by three data-dependent MS/MS spectra (m/z 100 – 2000, scan rate 20 Hz) for the three most intense ions per scan. The ions were selected when they reach an intensity threshold of 1000, isolated at the tuning file Q1 resolution, fragmented using collision induced dissociation (CID) with collision energy ramp (CE 20 – 50 eV), and excluded for 0.05 s (one MS scan) before being re-selected for fragmentation. The parameters used for the ESI source were: interface voltage 4 kV ( -3 kV for negative polarity), interface temperature 300 °C, nebulizing gas flow 3 L/min, and drying gas flow 10 L/min. The parameters used for the CDS probe were: interface voltage 4.5 kV (positive polarity) or -3.5 kV (negative polarity), and nebulizing gas flow 1 L/min.

In a separate analysis for more detailed fragmentation data, samples were prepared for injection as follows. Crude extracts were dried *in vacuo*, dissolved in $H_2O$ containing 0.5% formic acid, and loaded onto a Sep-Pak C18 column (Waters), which was activated with methanol. The column was washed twice with $H_2O$ containing 0.5% formic acid, then eluted with 80% ACN solution with 0.5% formic acid. Samples were dried *in vacuo*, and dissolved in $H_2O$:ACN:formic acid = 97:3:0.1. 100 ng of the extract was injected and analyzed by reverse-phase liquid chromatography on a nanoAcquity UPLC system (Waters) equipped with HSS-T3 C18 1.8 µm, 75 µm X 250 mm column (Waters). A gradient from 1% to 40% acetonitrile in 60 min was applied, [$Glu^1$]-fibrinopeptide B was used as lock mass compound and sampled every 30 s. Online MS/MS analysis was done using Synapt G2-Si HDMS mass spectrometer (Waters) with an $UDMS^E$ method set up as described[74]. Mass spectrum data were generated using ProteinLynx Global SERVER (PLGS, version 3.0.3), with $MS^E$ processing parameters with charge 2 lock mass 785.8426 Da.

*LC-MS based comparative metabolomics*

All raw data obtained from LC-MS analysis were converted to mzXML centroid files using Shimadzu LabSolutions Postrun Analysis. The converted files were imported and processed MZmine 2.5.3[75]. Throughout the analysis, *m/z* tolerance was set to 0.002 m/z or 10.0 ppm, RT tolerance was set to 0.05 min, noise level was set to 2.0E2 and minimum absolute intensity was set to 5.0E2 unless specified otherwise. Features were detected (polarity: positive, mass detector: centroid) and their chromatograms were built using the ADAP chromatogram builder[76] (minimum group size in number of scans: 10; group intensity threshold: 2.0E2). The detected peaks were smoothed (filter width: 9), and the chromatograms were deconvoluted (algorithm: local minimum search; Chromatographic threshold: 90%; search minimum in RT range: 0.05; minimum relative height: 1%; minimum ratio of peak top/edge: 2; peak duration 0.03 – 3.00 min). The detected peaks were deisotoped (maximum charge: 5; representative isotope: lowest *m/z*). Peak lists from different extracts were aligned (weight for RT = weight for *m/z*; compare isotopic pattern with a minimum score of 50%). Missing peaks detected in at least one of the sample were filled with the gap filling algorithm (RT tolerance: 0.1 min). Among the peaks, we identified fragments (maximum fragment peak height: 50%), adducts ($[M+Na]^+$, $[M+K]^+$, $[M+NH_4]$, maximum relative adduct peak height: 3000%) and complexes (Ionization method: $[M+H]^+$, maximum complex height: 50%). Duplicate peaks were filtered. Artifacts caused by detector ringing were removed (*m/z* tolerance: 1.0 m/z or 1000.0 ppm) and the results were filtered down to the retention time of interest. The aligned peaks were exported to a

MetaboAnalyst file. From here, peaks were additionally filtered to keep only peaks present in all three replicates, using in-house scripts. The resulting peak list was uploaded to MetaboAnalyst[77], log transformed and normalized with Pareto scaling without prior filtering. Missing values were filled with half of the minimum positive value in the original data. Heatmaps and volcano plots were generated using default parameters.

*Mass spectrometry-based quantitative proteomics*

20 uL of dense spore stocks were inoculated in NMMP and grown for 7 days as described above. 1 mL samples were taken after 2 and 7 days. Mycelium was gathered by centrifugation and washed with disruption buffer (100 mM Tris-HCl, pH 7.6, 0.1 M dithiothreitol). The samples were sonicated for 5 minutes (in cycles off 5s on, 5s off) to disrupt the cell wall, and centrifuged at max speed for 10 minutes to collect the proteins. Proteins were then precipitated using chloroform-methanol[78]. The dried proteins were dissolved in 0.1% RapiGest SF surfactant (Waters) at 95°C. Protein digestion steps were done according to van Rooden et al[74]. After digestion, formic acid was added for complete degradation and removal of RapiGest SF. Peptide solution containing 8 µg peptide was then cleaned and desalted using the STAGETipping technique[79]. Final peptide concentration was adjusted to 40 ng/µl with 3% acetonitrile, 0.5% formic acid solution. 200 ng of digested peptide was injected and analysed by reverse-phase liquid chromatography on a nanoAcquity UPLC system (Waters) equipped with HSS-T3 C18 1.8 µm, 75 µm X 250 mm column (Waters). A gradient from 1% to 40% acetonitrile in 110 min was applied, [$Glu^1$]-fibrinopeptide B was used as lock mass compound and sampled every 30 s. Online MS/MS analysis was done using Synapt G2-Si HDMS mass spectrometer (Waters) with an $UDMS^E$ method set up as described[74].

Mass spectrum data were generated using ProteinLynx Global SERVER (PLGS, version 3.0.3), with $MS^E$ processing parameters with charge 2 lock mass 785.8426 Da. Reference protein database was downloaded from GenBank with the accession number GCA_001278075.1. The resulting data were imported to ISOQuant[80] for label-free quantification. TOP3 quantification result from ISOQuant was used when further investigating the data.

*Iodoacetamide treatment*

Reaction mixtures were prepared based on earlier reported studies[66]. 20 µL reaction mixtures containing 0.25 mg/mL purified peptide, 13 mM TCEP, 25 mM IAA and 250 mM HEPES (pH = 8.0) in $H_2O$ were left at room temperature for 1 hour in the dark. Reaction mixtures were cleaned using the STAGETipping technique[79].

*DTT treatment*

Reaction mixtures were prepared based on earlier reported studies[65]. 20 µL reaction mixtures containing 0.25 mg/mL purified peptide, 10 mM DIPEA and 500 mM DTT in 1:1 $CHCl_3$ were left at room temperature for 18 hours. Mixtures were dried, dissolved in 0.5% formic acid in water and cleaned using the STAGETipping technique[79].

*Protein hydrolysis*

0.2 mg of purified peptide was dissolved in 3 mL 6M HCl and sealed inside a glass ampule, based on earlier studies[81]. The mixture was heated to 110°C for 24 hours. The HCl was removed by repeated drying and dissolving of the peptide with $H_2O$. The peptide was afterwards dissolved in 50 µL $H_2O$ and analyzed with LCMS as described above.

Bioinformatics

**decRiPPter pipeline**

*Genome data preparation*

As input, decRiPPter uses a set of genomes from species that are part of the same taxonomic group (e.g., genus, family), which it requires for its comparative genomic analyses. decRiPPter downloads genomes from NCBI[82] based on NCBI taxonomic identifiers of species, genera or higher orders of classification. Additional requirements for level of assembly (e.g. "Representative genome") can also be given. decRiPPter can reannotate genomes with prodigal 2.6.3[83], and automatically does so when DNA fasta files are given as input. In addition, users may analyze their own genomes, in isolation or in conjunction with downloaded genomes.

*SVM*

To predict RiPP gene clusters, we first collected positively and negatively labeled training data. The positive training data was collected from MIBiG[69] and recent literature, resulting in 175 RiPP precursors across ten classes (Supplemental Document 4). For the negative training set, we generated a set of 20,000 short non-precursor sequences. Half of these were randomly selected from a set of 35,000 short proteins (<175 amino acids long) from Uniprot (queried June 2014) that were not similar to RiPP precursors based on an NCBI blastp search. The other half were randomly selected from a set of 17,000 translated intergenic sequences between a stop codon and the next start codon of sizes 30-300 nt taken from 10 genomes across the bacterial tree of life: *Escherichia coli*, *Bacillus subtilis*, *Streptomyces coelicolor*, *Bacteroides fragilis*, *Rhizobium etli*, *Chloroflexus aurantiacus*, *Synechococcus* sp. PCC 7002, *Opitutus terrae*, *Acidobacterium capsulatum* and *Pirellula staleyi* (see Online Data at Zenodo). For all sequences from both the positive and negative training sets, we computed several physio-chemical properties, such as its length, hydrophobicity, charge, counts of canonical amino-acid residues and classes of amino acids, and highest counts of, e.g., cysteines and serines within contiguous blocks of 20 or 30 amino acids (See Online Data at Zenodo).

We then utilized Scikit-Learn implementations of several different supervised machine-learning algorithms. We varied many parameters associated with a given algorithm (e.g., different kernel functions, a range of different values for penalty parameters, different penalty functions, etc.). Furthermore, we mapped the accuracy as a function of scaling the dataset or changing class weights to take into account the unbalanced dataset (only ~1% of gene clusters in our dataset represent known RiPPs). The RiPP cluster classification accuracy of each combination of scaling, algorithm, and the corresponding set of parameters was evaluated using accuracy and area under receiver operating characteristics (ROC) curve, and leave-one-class-out cross-validation. The SVM algorithm with cubic kernel function, scaling, and penalty parameter C, kernel coefficient gamma, and independent term in kernel function coef0 of 0.158, 0.333, 2.154, respectively, was found to perform most accurately and was used for further analysis.

*COG scores calculation*

To calculate the relative frequency of occurrence of each gene, we constructed a pipeline to find all groups of homologous genes (figure S1). In the first step, protein-coding genes for which orthology can confidently be assigned are grouped into Clusters of Orthologous Groups (COGs). All proteins are aligned to one another using DIAMOND[34], and all bidirectional best hits (BBHs) are identified that share at least 60.0% similarity (figure S1A). We established two requirements for genes to be confidently annotated as orthologs, based on recent papers[35,36]: 1) they should constitute BBHs, and 2) their immediate genomic surroundings should be conserved, i.e. the two flanking genes should also be bidirectional best hits between the two genomes. Genes fulfilling these two criteria are paired together, resulting in groups of orthologous genes. Among these groups, decRiPPter then selects those that are completely conserved across all genomes: each group should contain at least one ortholog in each genome, and all orthologs in the group should all fulfill the same requirements for each genome pair. These groups are considered true Clusters of Orthologous Genes (trueCOGs; figure S1B).

In the second step, a cutoff for protein-coding gene sequence identity is determined for each genome pair, in order to separate orthologs as well as recently evolved paralogs from more distantly related homologs. For any given pair of genomes, the distribution of sequence identities of all gene pairs of their trueCOGs is calculated. The cutoff is then calculated as the average percentage identity, minus three times the standard deviation (figure S1C). Any two aligned genes with a percentage identity higher than this cutoff are considered to be functionally closely related to one another and paired up. The resulting groups of homologous genes were clustered with the Markov Cluster Algorithm[37] (figure S1D). From these groups, the relative frequency of occurrence of groups of homologous genes across all query genomes is calculated, called the COG-score (figure S1E).

In cases when insufficient numbers of trueCOGs (<= 10) could be found in our analyses (because the set of genomes was too diverse, and/or contained too many draft genomes that each miss some of the trueCOGs), the genomes were rearranged into smaller subgroups. We used two general rules to create the groups: 1) Groups should be as large as possible, so that trueCOGs found are conserved across many species, and represent conserved widespread genes. 2) Genomes should be compared to as many other genomes as possible, so as not to introduce bias into the calculation of the COG-score. To fulfil both requirements, partially overlapping subgroups were formed, with the goal of letting each genome be a part of a collection of subgroups that together covered as many of the genomes as possible. To form the subgroups, a pair of genomes with the highest number of trueCOGs was used as a seed, and genomes were added one at a time until the number of trueCOGs dropped below the set cutoff. All the genomes in the group were said to be linked together by this group. The process of group formation was then repeated, starting with genomes for which no group had yet been formed. If all genomes were already part of at least one subgroup, the genomes were selected which were linked to the fewest genomes via the groups they were part of. The process was terminated when adding additional groups did not increase the number of links between genomes for several successive iterations.

*Gene cluster formation*

In this stage, decRiPPter identifies putative operon-like gene clusters around each candidate precursor peptide-encoding gene, by either of two different methods: In the first method, called the simple method, genes in the same strand orientation as the candidate precursor peptide-encoding gene are added to the putative gene cluster if the intergenic distance to the previous gene is within a given cutoff. The second method, called the island method, uses both intergenic distance and levels of conservation (COG-score) to determine the gene clusters. First, all genes in the same strand orientation within 750 nucleotides of one another are identified and then grouped into islands. Within islands, genes should be almost directly adjacent (intergenic distance: <= 50 nucleotides). We then fused the islands together using the COG-scores (see above), building on the assumption that genes in a gene cluster should all have similar levels of conservation. Islands were fused together if the average of their COG-scores was within a set range (0.1 plus the sum of the standard deviations of both islands). Not all gene families have similar COG scores when they occur within the gene clusters thus formed; e.g., genes encoding ABC-transporters frequently have close relatives in other biomolecular systems and therefore often have higher COG scores. Hence, to counteract gene cluster formation breaking off prematurely, up to two outlier genes are allowed when fusing islands, if, after adding the outliers, more islands can be added that are within the range for COG-score deviation. Intergenic distances and cutoffs were iteratively finetuned to ensure gene clusters in known RiPP BGCs would be effectively found. Finally, gene clusters that overlap or lie within 50 nucleotides of one another are fused together.

*Annotation*

For purposes of data exploration (annotation and visualization), each gene cluster is extended to include the 5 flanking genes on either side, and all encoded proteins in the extended gene clusters are annotated with Pfam 31.0[28] and TIGRFAM[29]. Lists were compiled of all TIGRFAM and Pfam domains associated with either peptidases, transporters, regulators, using a combination of keyword searches on the Pfam and TIGRFAM websites, combined with manual curation. A list of protein domains associated with biosynthetic activity was constructed by linking Pfam domains to E.C. numbers, either through the KEGG or through the GO database. Biosynthetic TIGRFAM domains were taken directly from the database. Each domain linked to an E.C. number was assumed to have enzymatic activity. The biosynthetic domain list was further expanded with domains used in the ClusterFinder[27] algorithm that were indicative of a biosynthetic gene cluster. The resulting lists are used by decRiPPter to mark proteins either as a regulator, peptidase, transporter or biosynthetic enzyme, in that order, by seeing if any of the identified domains overlapped with the domains in the precompiled lists.

*Clustering*

To cluster the detected gene clusters, the distance between them is calculated in two different ways: 1) amino acid sequences of candidate precursor peptide-encoding genes in the gene clusters are aligned with NCBI BLAST[84] blastp (cutoff: 30 bitscore), and 2) the content of the gene clusters is compared by calculating the Jaccard index of their constituent protein domains (cutoff: 0.5). Gene clusters are paired only if they are paired by both methods. The distance between paired gene clusters is calculated as the average between the Jaccard index and the percentage identity of the aligned precursors. Finally, pairs are clustered using MCL.

*Overlap with antiSMASH*

Overlap with antiSMASH was determined using antiSMASH 4.0[42] on minimal mode.

*Availability*

The decRiPPter pipeline is available at https://github.com/Alexamk/decRiPPter/.

**Data analysis**

*Streptomyces analysis*

For genome mining of *Streptomyces*, we downloaded all available genomes falling under the taxonomic identifier for *Streptomyces* (1883). Genomes were reannotated with prodigal 2.6.3, and processed with the pipeline described above. Gene clusters were formed with the island method.

The results of the strict and the mild filter is available at http://www.bioinformatics.nl/~medem005/decRiPPter_strict/index.html and http://www.bioinformatics.nl/~medem005/decRiPPter_mild/index.html, respectively.

*Genomic context analysis*

CORASON[85] was used with the number of flanking genes set to 15, on the *Streptomyces* genomes analyzed with the query of interest. Results were parsed using in-house scripts and compared to decRiPPter output. NCBI BLAST was used to find additional homologs of genes of interest within the clusters, with a cutoff of 30 percent ID similarity.

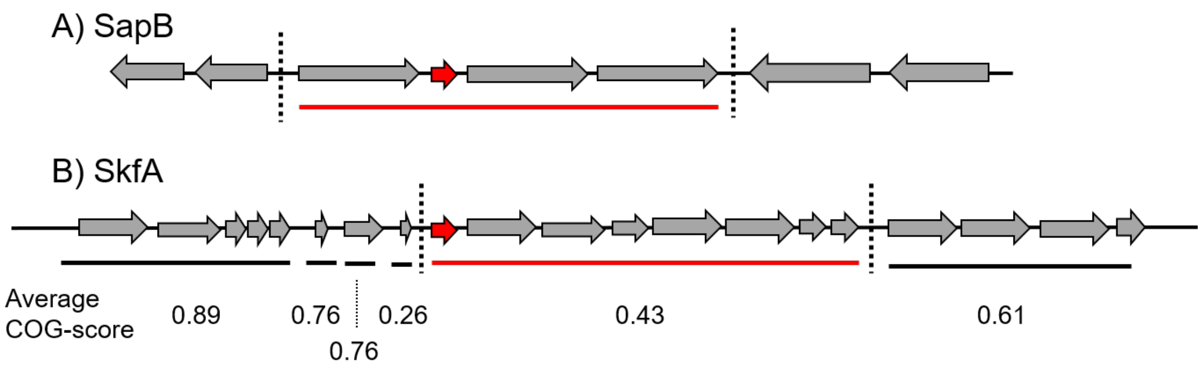

**Figure S1. decRiPPter forms putative gene clusters around candidate precursor peptide-encoding genes.** Two examples are provided here to illustrate identification of putative gene clusters in decRiPPter. A) In the *sapB* gene cluster, four genes form the main BGC. These four genes are sequential, share the same strand orientation and lie within a small distance of one another (<= 50 nt). They are therefore fused together into a single gene cluster. The flanking genes are on opposite strands, and therefore not considered. B) The *skfA* BGC consists of eight genes sequential genes that share the same strand orientation. However, it is flanked by several other genes that also share the same strand orientation, within relatively short intergenic distances (<= 200 nucleotides). Using the island method, the genes are first fused into six islands, within 50 nucleotides distance of one another (indicated by lines underneath the genes). These islands may then be fused depending on the COG-score, which does not happen here because the difference is too large. The result is that the flanking genes, with a too high COG-score, are removed, and the correct BGC remains.

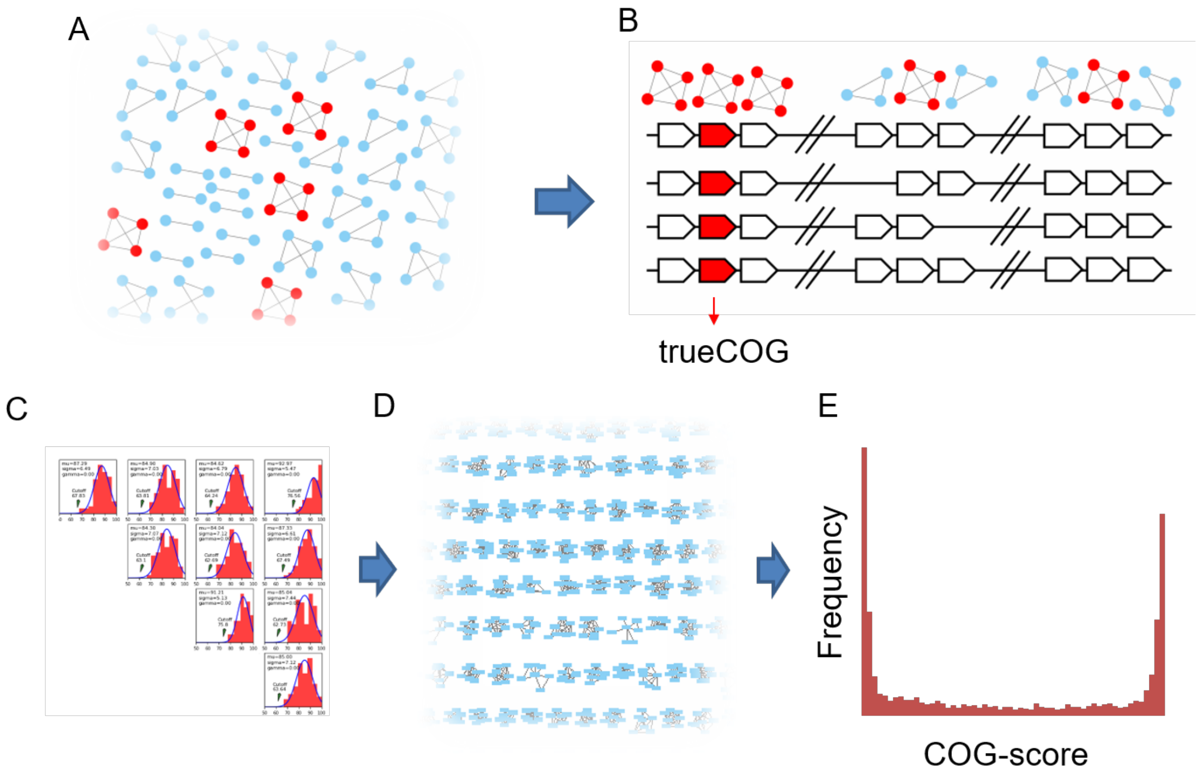

**Figure S2**. **decRiPPter determines the frequencies of occurrence of genes to calculate the COG score.** In this example, the COG score of four genomes is calculated. A) All encoded proteins are aligned to find bidirectional best hits (BBHs; edges). All clusters of BBHs conserved across all genomes are displayed as red. If one genome does not contain a homologous gene, or the gene in question is not a BBH with all genes from the cluster from other genomes, it is not considered a conserved group of BBHs. B) If the flanking genes of the clusters of BBHs are also part of clusters of BBHs, the center genes are considered to form a true Cluster of Orthologous Genes (trueCOG). Of the three cases displayed here, only the leftmost group passes this criterion; for the center group, not all genes are conserved, and for the right group, not all genes are BBHs with one another in the flanking groups. C) The genes in all trueCOGs between each genome pair are used to create a sequence identity cutoff to use for all protein-coding genes in a given pair of genomes. D) All genes are paired using the sequence identity cutoffs determined in the previous step. E) The COG-score is calculated for each gene. Typically, a bimodal distribution can be seen, with many genes either conserved across all genomes, or only present in a single organism.

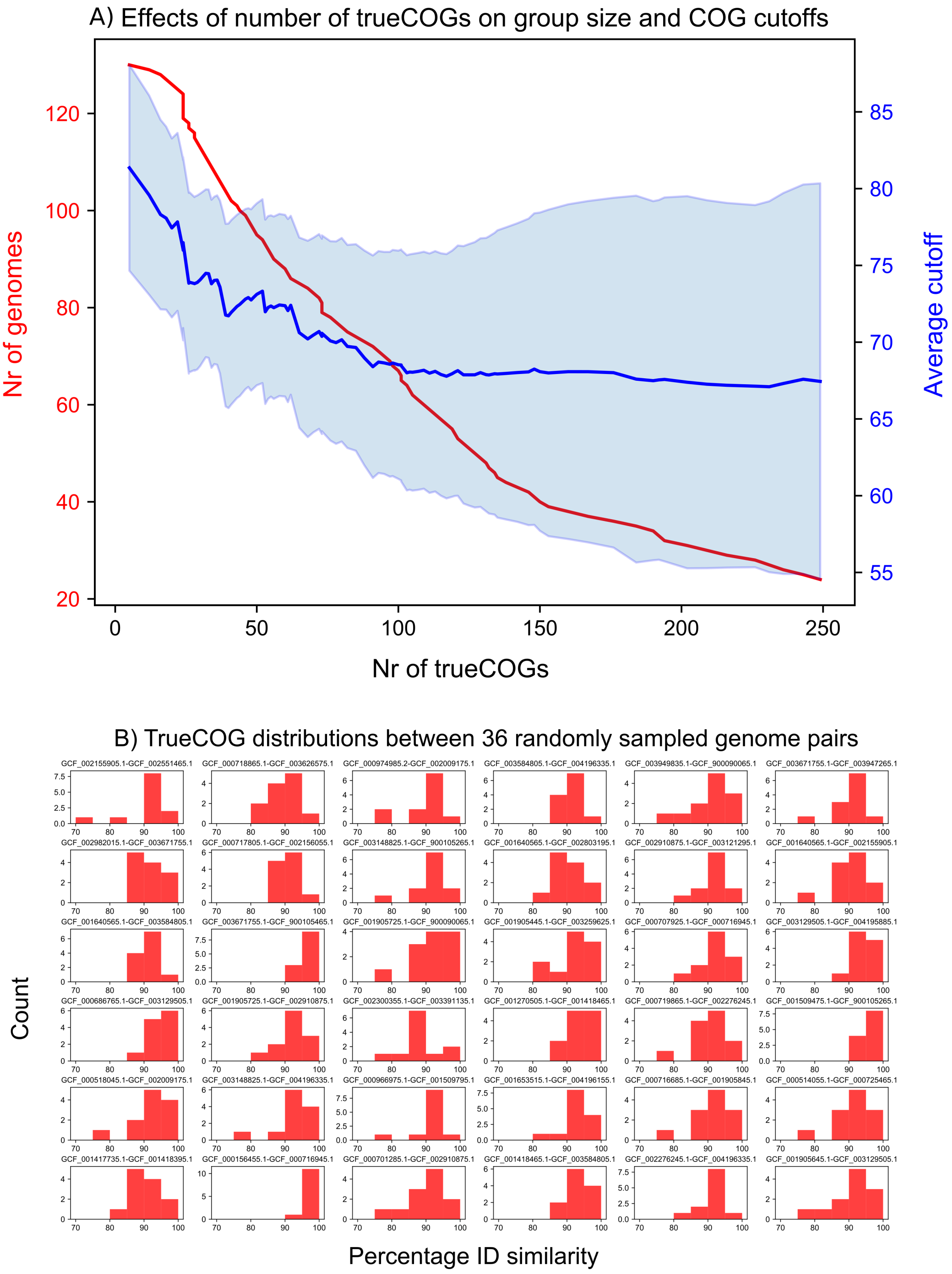

**Figure S3.** A) As the minimum number of trueCOGs increases, the number of genomes that can be analyzed together (red line) decreases. In addition, the average COG cutoff (blue line) decreases when more trueCOGs are added, and the spread of COG cutoffs (shaded area; average cutoff +- the standard deviation) increases, suggesting that additional trueCOGs that were added were less conserved and showed higher variability in sequence similarity. B) TrueCOG distribution between 36 randomly sampled genome pairs. Based on these distributions, COG cutoffs were determined.

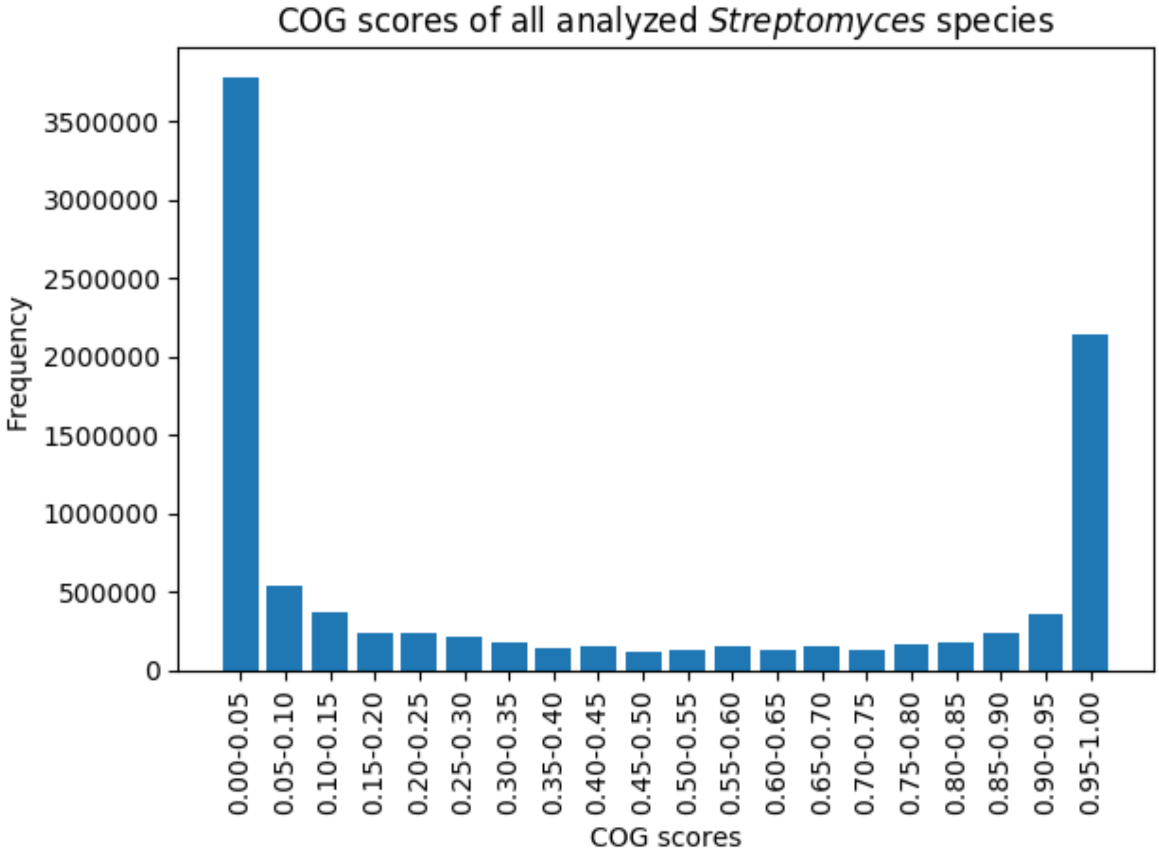

**Figure S4. COG scores of all genes in all 1,295 analyzed *Streptomyces* genomes**. A high COG score indicates presence of homologs in many different genomes, while a low COG score indicates a more infrequent distribution. COG scores were calculated as described in the methods.

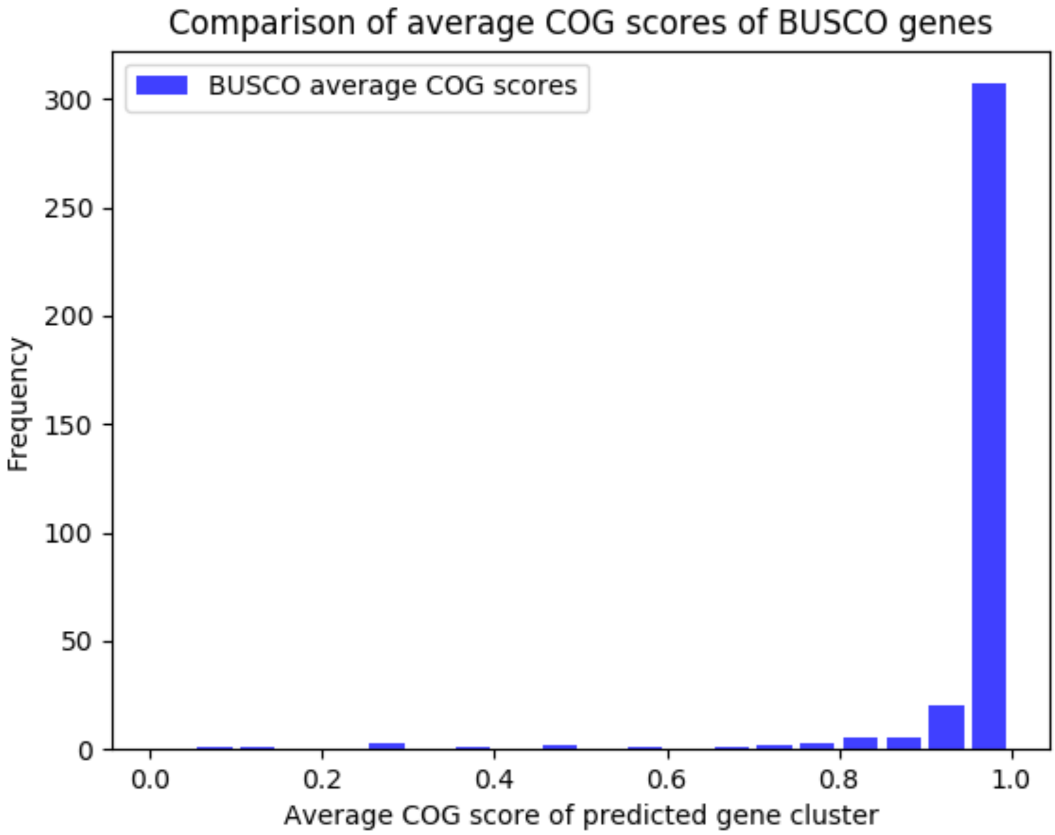

**Figure S5**. **Comparison of average COG scores of BUSCO genes**. The average of each BUSCO[26] gene was calculated for each genome analyzed.

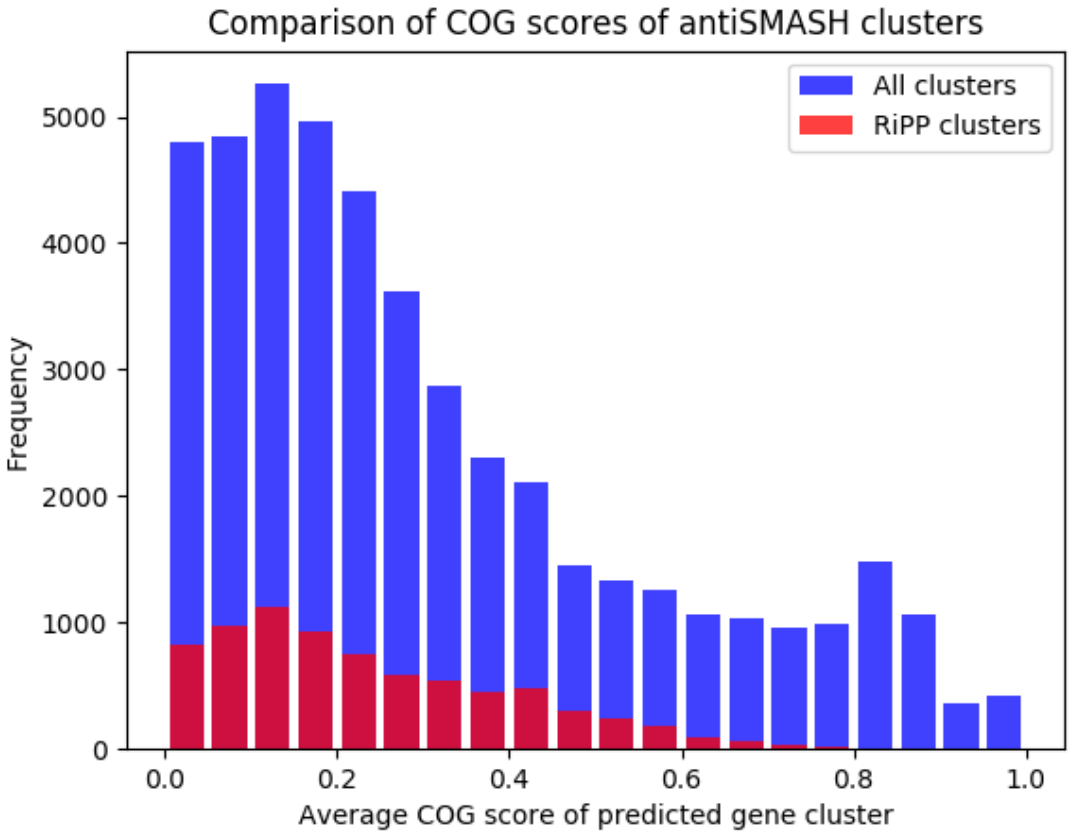

**Figure S6. Comparison of COG scores of antiSMASH-detected gene clusters**. COG scores were averaged over all genes in the predicted gene clusters. COG scores averaged 0.311 +- 0.249 for all gene clusters, and 0.234 +- 0.166 for RiPP gene clusters.

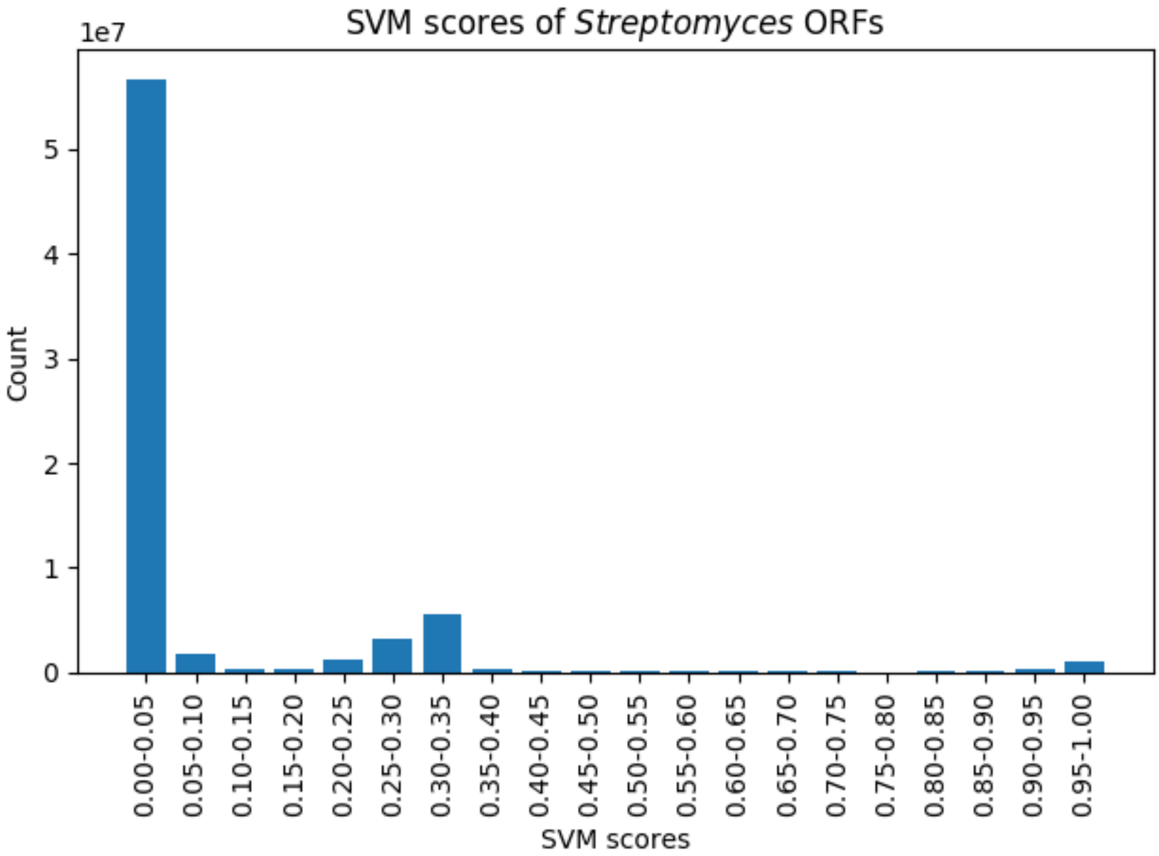

**Figure S7. Distribution of the scores assigned by the SVM classifier**. These scores indicate the likelihood that a specific gene encodes for a precursor. A total of $7,1 * 10^7$ small open reading frames were analyzed. Based on the slight enrichment of precursors with scores >= 0.90, the cutoff was set at 0.9 (see main text).

z

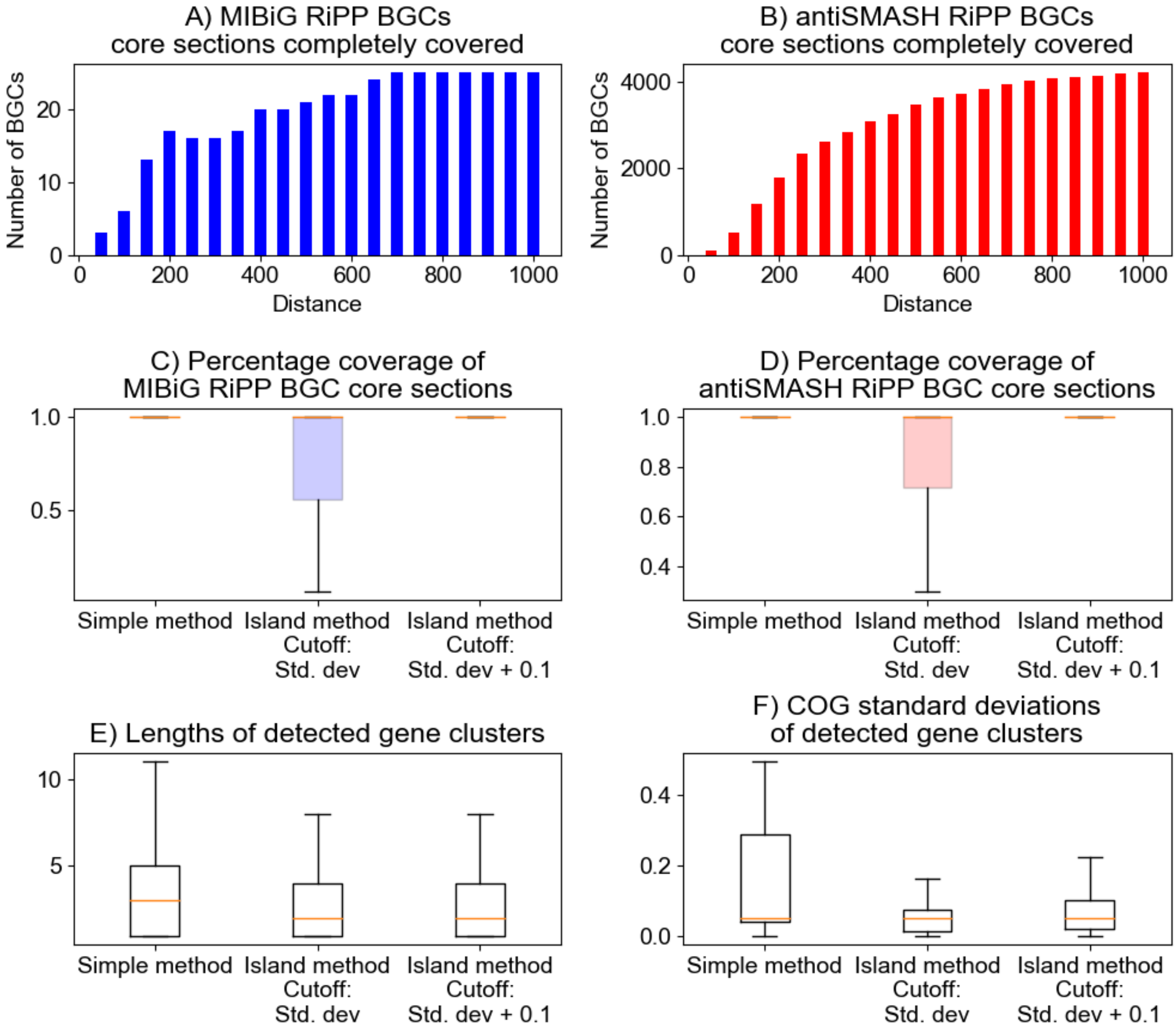

**Figure S8. Gene cluster formation effectively covers antiSMASH and MIBiG BGC core gene sections.** In the simple gene cluster formation method, genes are sequentially added as long as they are in the same strand orientation, within a certain distance. At a distance of 700 nucleotides, all MIBiG core gene sections are covered (A), as well as 91% (3947/4321) of antiSMASH core gene sections. (B). In the 'island method', genes are first fused into islands, which may be further fused if their average COG-scores are within a cutoff. Using just the standard deviation of the islands as a cutoff resulted in incomplete coverage of both the MIBiG and the antiSMASH core sections (C, D, middle boxes). Increasing the cutoff to the standard deviation plus 0.1 resulted in comparable coverage (C, D, right boxes) of these sections when compared to the simple method (C, D, left boxes). In addition, the overall gene cluster length (E) and variation of COG scores (F) within all formed gene clusters decreased.

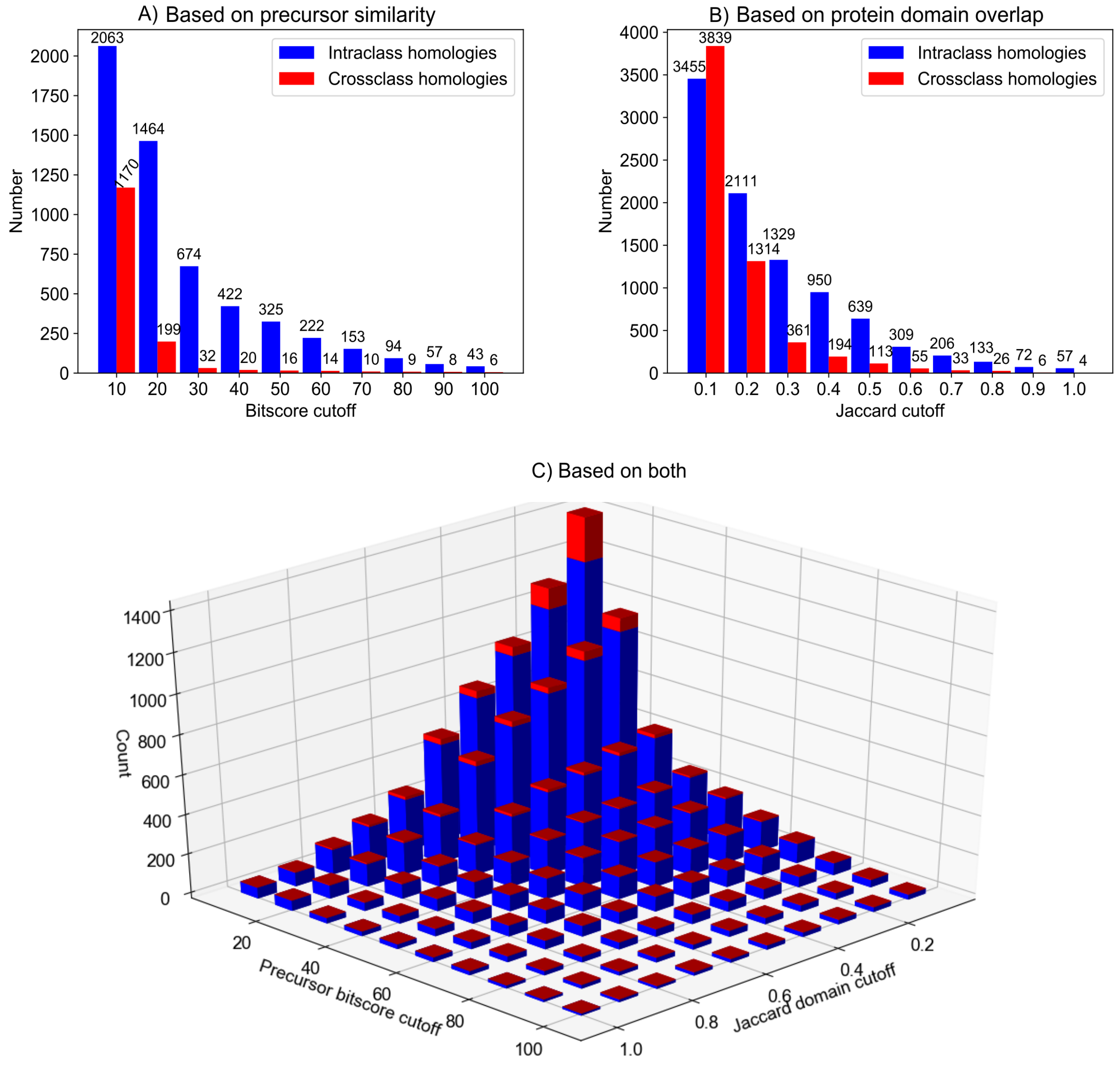

**Figure S9. Combining precursor similarity with domain similarity is an effective strategy to group RiPP subclasses.** Starting at precursor similarity bitscore cutoffs of 20 and Jaccard scores of overlapping protein domains found in MIBiG RiPP BGCs of 0.4, the number of intraclass homologies is larger than the number of crossclass homologies. Combining the two methods greatly decreases the number of cross-class homologies found, proving it as an effective method to group RiPP BGCs of different subtypes.

**Figure S10. Alignment of precursors belonging to the characterized family of type V lanthipeptides.** Precursors were aligned with MUSCLE[27] and visualized with BoxShade.

```
prod_559746     1 ---MHTM-TETDLLSGYTAYTTAEELDQFDGKAAPAATTPVLAPILI-----RASIIAARSSQQCA----AGIAAAGGGIWRTIRKVC
prod_4312120    1 ---MQNV-TEKDLFDGYTAYTSAEELGLHDGATAGPAFSPTV-PWAI-----QATVISARSSQACA----AALGSLAA---KTVEKKC
prod_4312121    1 ---MQNV-TEKDLFDGYTAYTSAEELGLHDGATAGPAFSPTV-PWAI-----QATVISARSSQACA----AALGSLAA---KTVEKKC
prod_9638834    1 ---MQNV-TEQDLFDGYTAYTSAEELGLHDGKDAAPAFSPTI-PWAI-----RATIISARSSQQCA----AALGSLAA---KTVENKC
prod_1888002    1 ---MQNV-TEKDLFDGYTAYTSAEELGLHDGQEAAPAFSPTI-PWAI-----RATIITARSSQQCA----AALGSLAA---KTVENKC
prod_1892473    1 ---MQNV-TEKDLFDGYTAYTSAEELGLHDGQEAAPAFSPTI-PWAI-----RATIITARSSQQCA----AALGSLAA---KTVENKC
prod_1898975    1 ---MQNV-TEKDLFDGYTAYTSAEELGLHDGQEAAPAFSPTI-PWAI-----RATIITARSSQQCA----AALGSLAA---KTVENKC
prod_2702012    1 ---MQNV-TEKDLFDGYTAYTSAEELGLHDGQEAAPAFSPTI-PWAI-----RATIITARSSQQCA----AALGSLAA---KTVENKC
prod_4125916    1 ---MQNV-TEKDLFDGYTAYTSAEELGLHDGQEAAPAFSPTI-PWAI-----RATIITARSSQQCA----AALGSLAA---KTVENKC
prod_4204099    1 ---MQNV-TEKDLFDGYTAYTSAEELGLHDGQEAAPAFSPTI-PWAI-----RATIITARSSQQCA----AALGSLAA---KTVENKC
prod_5620390    1 ---MQNV-TEKDLFDGYTAYTSAEELGLHDGQEAAPAFSPTI-PWAI-----RATIITARSSQQCA----AALGSLAA---KTVENKC
prod_5701534    1 ---MQNV-TEKDLFDGYTAYTSAEELGLHDGQEAAPAFSPTI-PWAI-----RATIITARSSQQCA----AALGSLAA---KTVENKC
prod_5937191    1 ---MQNV-TEKDLFDGYTAYTSAEELGLHDGQEAAPAFSPTI-PWAI-----RATIITARSSQQCA----AALGSLAA---KTVENKC
prod_6249001    1 ---MQNV-TEKDLFDGYTAYTSAEELGLHDGQEAAPAFSPTI-PWAI-----RATIITARSSQQCA----AALGSLAA---KTVENKC
prod_6819619    1 ---MQNV-TEKDLFDGYTAYTSAEELGLHDGQEAAPAFSPTI-PWAI-----RATIITARSSQQCA----AALGSLAA---KTVENKC
prod_710895     1 ---MQNV-TEKDLFDGYTAYTSAEELGLHDGQEAAPAFSPTI-PWAI-----RATIITARSSQQCA----AALGSLAA---KTVENKC
prod_7443641    1 ---MQNV-TEKDLFDGYTAYTSAEELGLHDGQEAAPAFSPTI-PWAI-----RATIITARSSQQCA----AALGSLAA---KTVENKC
prod_7703323    1 ---MQNV-TEKDLFDGYTAYTSAEELGLHDGQEAAPAFSPTI-PWAI-----RATIITARSSQQCA----AALGSLAA---KTVENKC
prod_8242019    1 ---MQNV-TEKDLFDGYTAYTSAEELGLHDGQEAAPAFSPTI-PWAI-----RATIITARSSQQCA----AALGSLAA---KTVENKC
prod_8466597    1 ---MQNV-TEKDLFDGYTAYTSAEELGLHDGQEAAPAFSPTI-PWAI-----RATIITARSSQQCA----AALGSLAA---KTVENKC
prod_8698113    1 ---MQNV-TEKDLFDGYTAYTSAEELGLHDGQEAAPAFSPTI-PWAI-----RATIITARSSQQCA----AALGSLAA---KTVENKC
prod_8721923    1 ---MQNV-TEKDLFDGYTAYTSAEELGLHDGQEAAPAFSPTI-PWAI-----RATIITARSSQQCA----AALGSLAA---KTVENKC
prod_8902069    1 ---MQNV-TEKDLFDGYTAYTSAEELGLHDGQEAAPAFSPTI-PWAI-----RATIITARSSQQCA----AALGSLAA---KTVENKC
prod_9724047    1 ---MQNV-TEKDLFDGYTAYTSAEELGLHDGQEAAPAFSPTI-PWAI-----RATIITARSSQQCA----AALGSLAA---KTVENKC
prod_1317692    1 ---MQNV-TEKDLFDGYTAYTSAEELGLHDGKEAAPAFSPTI-PWAI-----RATIISARSSQQCA----AALGSLAA---KTVENKC
prod_3048582    1 ---MQNV-TEKDLFDGYTAYTSAEELGLHDGKEAAPAFSPTI-PWAI-----RATIITARSSQQCA----AALGSLAA---KTVENKC
prod_398364     1 ---MQSTQNEKDLFEGYTAYTSAEELGLYDGKDAAPAFSPTI-PWAI-----RATIITARSSQQCA----AAIGSLTA---KTIENKC
prod_7467458    1 ---MQSTQNEKDLFEGYTAYTSAEELGLYDGKDAAPAFSPTI-PWAI-----RATIITARSSQQCA----AAIGSLTA---KTIENKC
prod_5042396    1 ---MQNV-NEKDLFDGYTAYTSAEELGLYDGKDAAPAFSPTI-PWAI-----RAGLITARSSQQCA----AAIGSFTA---RTIESKC
prod_1644796    1 ---MNAS---AHLIAGYTAYTTAAEFDA-SITADAPAVTPAT-P--------SIALSIAESSYACG----AGVGASIG---ITFTKGC
prod_7595003    1 ---MNAS---AHLIAGYTAYTTAAEFDA-SITADAPAVTPAT-P--------SIALSIAESSYACG----AGVGASIG---ITFTKGC
prod_9224211    1 ---VNTT---ENLIAGYTAYTSAQEIEA-THAEEAPGATPSV---------LSFIATSGWA---CG----AGIGTSIG---VTAAKGC
prod_4694754    1 ---VNTT---DTLLAGYAAYTSADEIAA-AQDGGAPEISPVS----L-----SIAVSIAESSYACS----AGLSMSVG---VTVGKGC
prod_4694755    1 ---VNTT---DTLLAGYAAYTSADEIAA-AQDGGAPEISPVS----L-----SIAVSIAESSYACS----AGLSMSVG---VTVGKGC
prod_7200544    1 ---MNTS---DNLMAGYATYTSADEIAA-TLDGGAPEISPVS----L-----SIAVSITESSYACG----AGISLSVG---WTVGKGC
prod_9224208    1 ---MNTA---DQLMAGYAVYTTSDEIGA-GAAADAPAISPVS---IF-----SAASSVECAIFSAG----VVTSASAG---GTVAGNC
prod_4694758    1 ---MNTA---DQLIAGYTAYTDSAEIAA-DATAEAPAISPTT----------TITIVSVESVLASI----GASASFSAG--YTVSSGC
prod_7200547    1 ---MNNT---DQLIAGYTAYTDSAEIAA-DASAEAPAITPTT----------TITIVSVESAVFSI----GGAASFSAG--YTISSGC
prod_326225     1 --MSHDQNTLEELVTGYESYADADEIEV-DAVTGAPATTPFCGA--------AASFMLSYV---------------------TTNGPG
prod_326226     1 --MTNDQSTLEDLVTGYESYADADEIEV-DAVTGAPATTPFCGA--------VASFALSYV---------------------TTNGPG
prod_6174086    1 ---VKTQ----DLIAGYAAYVDVAELNV-SAASEAPATSPVCFAAATSSAACLAATSSGWCVAGAG----AGVGGGIA---QSVKHGC
prod_6174087    1 ---M----ELDEMISGYDTYVDVAELDV-SAQSEAPATSPTC-----------FIASVGLS--------------------YQITKDL
prod_9167739    1 ---MQNDIEIMELVGGFEAYTEAAELNM-EASVEAPAATPTA------------TIVYTKFS----------VASVT----LTAKKGC
prod_2743547    1 VQKNDTV-DIMELVGGFEAYAEAAELNF-EASADAPAITPTL-----------TTIAYTKVS----------VASVSA----SIKVGC
prod_5868070    1 VQNIENV-EIMELVGGFEAYAQAAELNF-EASADAPAITPTL-----------TTIAYTKVTVAGT----A----------ASIKWTC
prod_1221493    1 ---MDTH----ELIEGFDAYVEAEELNE-DAMVDAPATTVPC-----------------------------TVASF-----ATGYFSC
prod_3289459    1 ---MDTH----ELIEGFDAYVEAEELNE-DAMVDAPATTVPC-----------------------------TVASF-----ATGYFSC
prod_467494     1 ---MDTH----ELIEGFDAYVEAEELNE-DAMVDAPATTVPC-----------------------------TVASF-----ATGYFSC
prod_1100745    1 ---MEKATSIVELLSGYEAYSSVEEINL-SAASDAPATTWGCAA---------VSASISWM-----------SGQVVS---KTVDDGC
prod_2725163    1 ---MEKATSIVELLSGYEAYSSVEEINL-SAASDAPATTWGCAA---------VSASISWM-----------SGQVVS---KTVDDGC
prod_3616888    1 ---MEKATSIVELLSGYEAYSSVEEINL-SAASDAPATTWGCAA---------VSASISWM-----------SGQVVS---KTVDDGC
prod_5244387    1 ---MEKATSIVELLSGYEAYSSVEEINL-SAASDAPATTWGCAA---------VSASISWM-----------SGQVVS---KTVDDGC
prod_6023772    1 ---MEKATSIVELLSGYEAYSSVEEINL-SAASDAPATTWGCAA---------VSASISWM-----------SGQVVS---KTVDDGC
prod_6473304    1 ---MEKATSIVELLSGYEAYSSVEEINL-SAASDAPATTWGCAA---------VSASISWM-----------SGQVVS---KTVDDGC
prod_6857183    1 ---MEKATSIVELLSGYEAYSSVEEINL-SAASDAPATTWGCAA---------VSASISWM-----------SGQVVS---KTVDDGC
prod_8409183    1 ---MEKATSIVELLSGYEAYSSVEEINL-SAASDAPATTWGCAA---------VSASISWM-----------SGQVVS---KTVDDGC
prod_9246364    1 ---MEKATSIVELLSGYEAYSSVEEINL-SAASDAPATTWGCAA---------VSASISWM-----------SGQVVS---KTVDDGC
prod_9674514    1 ---MEKATSIVELLSGYEAYSSVEEINL-SAASDAPATTWGCAA---------VSASISWM-----------SGQVVS---KTVDDGC
prod_1949441    1 ---MEKATSIVELLSGYEAYSSAEEINL-SAATDAPATTWGCAA---------VSASVSWM-----------SGQVVS---KTVDDGC
prod_5478099    1 ---MEKATSIVELLSGYEAYSSAEEINL-SAATDAPATTWGCAA---------VSASVSWM-----------SGQVVS---KTVDDGC
prod_326224     1 ---MDNA-SMMDLVAGYNTYAEASELGI-QAVADAPATTPVCAATIA-----ASAVSSGWC---AS----AAASAAGG---ATYKLGC
sprA3           1 ---MQNNTEIMDLIANYDAYADVDELNV-TAAADAPATTPVCAA---------SVASSTW----CA----SAASAISG---ATYEAGC
prod_8036387    1 --MSNKSTVIADLVAGYDAYTEVDELNV-SAAAGAPATTWVC----------VSVVASRASSVKCGAWASAGASAVSG---ATYEITC
prod_2805062    1 --MDNKSTVITDLVAGYSTYTEAGELNV-SAAAGAPATTYIC----------ASVAISRVSSPRCA----ASASAVSG---ATYEWTC
prod_297319     1 --MDNKSTVITDLVAGYSTYTEAGELNV-SAAAGAPATTYIC----------ASVAISRVSSPRCA----ASASAVSG---ATYEWTC
prod_5421071    1 --MDNKSTVITDLVAGYSTYTEAGELNV-SAAAGAPATTYIC----------ASVAISRVSSPRCA----ASASAVSG---ATYEWTC
prod_5527162    1 --MDNKSTVITDLVAGYSTYTEAGELNV-SAAAGAPATTYIC----------ASVAISRVSSPRCA----ASASAVSG---ATYEWTC
prod_6582107    1 --MDNKSTVITDLVAGYSTYTEAGELNV-SAAAGAPATTYIC----------ASVAISRVSSPRCA----ASASAVSG---ATYEWTC
prod_8403914    1 --MDNKSTVITDLVAGYSTYTEAGELNV-SAAAGAPATTYIC----------ASVAISRVSSPRCA----ASASAVSG---ATYEWTC
prod_9151868    1 --MDNKSAVITDLVAGYSTYTEAGELNV-SAAAGAPATTYIC----------ASVAISRTSSVKCS----AAASAISG---ATYEWTC
prod_9381790    1 --MDNKSAVITDLVAGYSTYTEAGELNV-SAAAGAPATTYIC----------ASVAISRTSSVKCS----AAASAISG---ATYEWTC
prod_8036386    1 ---MKTT-TIMELAAGYDAYTGAEELEV-GATAEAPASTPLCAA--------AASAGVSWM-----------ASQFSA---RTISGGC
prod_9151867    1 ---MKTT-AIMELVAGYEVYADSAELQV-DATVNAPASTPAC-----------GAATVSWI-----------VSQFSA---KTVKDGC
prod_9381789    1 ---MKTT-AIMELVAGYEVYADSAELQV-DATVNAPASTPAC-----------GAATVSWI-----------VSQFSA---KTVKDGC
prod_2805061    1 ---MKTT-AIMELVAGYEVYADSAELQV-DAAVDAPASTPAC-----------AAATLSWI-----------VSQFSG---KTVKDGC
prod_297320     1 ---MKTT-AIMELVAGYEVYADSAELQV-DAAVDAPASTPAC-----------AAATLSWI-----------VSQFSG---KTVKDGC
prod_5421070    1 ---MKTT-AIMELVAGYEVYADSAELQV-DAAVDAPASTPAC-----------AAATLSWI-----------VSQFSG---KTVKDGC
prod_5527161    1 ---MKTT-AIMELVAGYEVYADSAELQV-DAAVDAPASTPAC-----------AAATLSWI-----------VSQFSG---KTVKDGC
prod_6582106    1 ---MKTT-AIMELVAGYEVYADSAELQV-DAAVDAPASTPAC-----------AAATLSWI-----------VSQFSG---KTVKDGC
prod_8403913    1 ---MKTT-AIMELVAGYEVYADSAELQV-DATVDAPASTPAC-----------AAATLSWI-----------VSQFSG---KTVKDGC
sprA1           1 MADLQQTGSISELVAGYDTYSEAGELVA-EAAADAPASTPTC-----------AAATISWL-----------GSQLTV---KTYKEGC
sprA2           1 ---MDKTGAITELIEGYDSYSDAEELNS-TAAAEAPATSAPC-----------GAASVSWL-----------ASQFTV---KTYKEGC
```

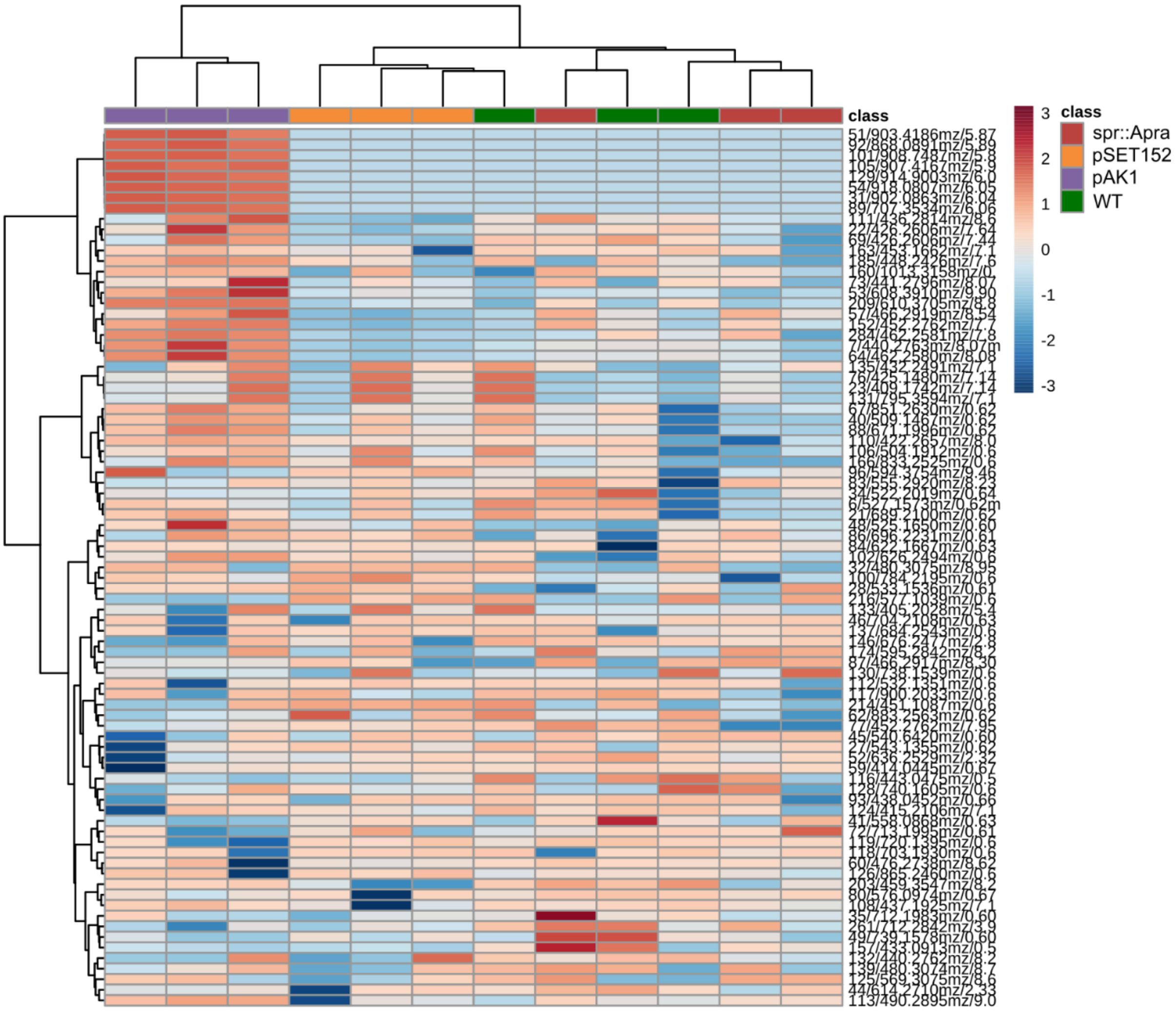

**Figure S11. Heatmap of extracted peaks reveals seven peaks that are uniquely observed in strains containing the expression construct pAK1.** Color of the area indicates a log-fold increase.

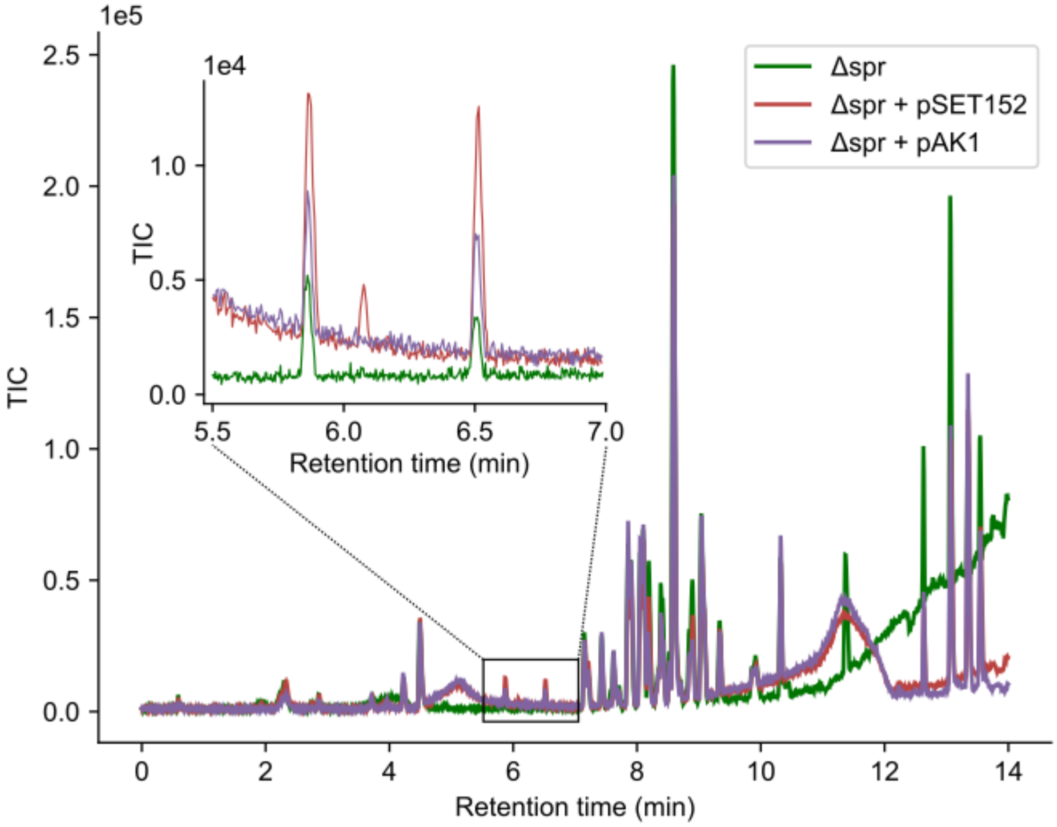

**Figure S12. Strains lacking the *spr* gene cluster are unable to produce the extracted products.** Removing the entire gene cluster from the genome nullified the effect of transforming pAK1 on the extracted masses (see figure 3).

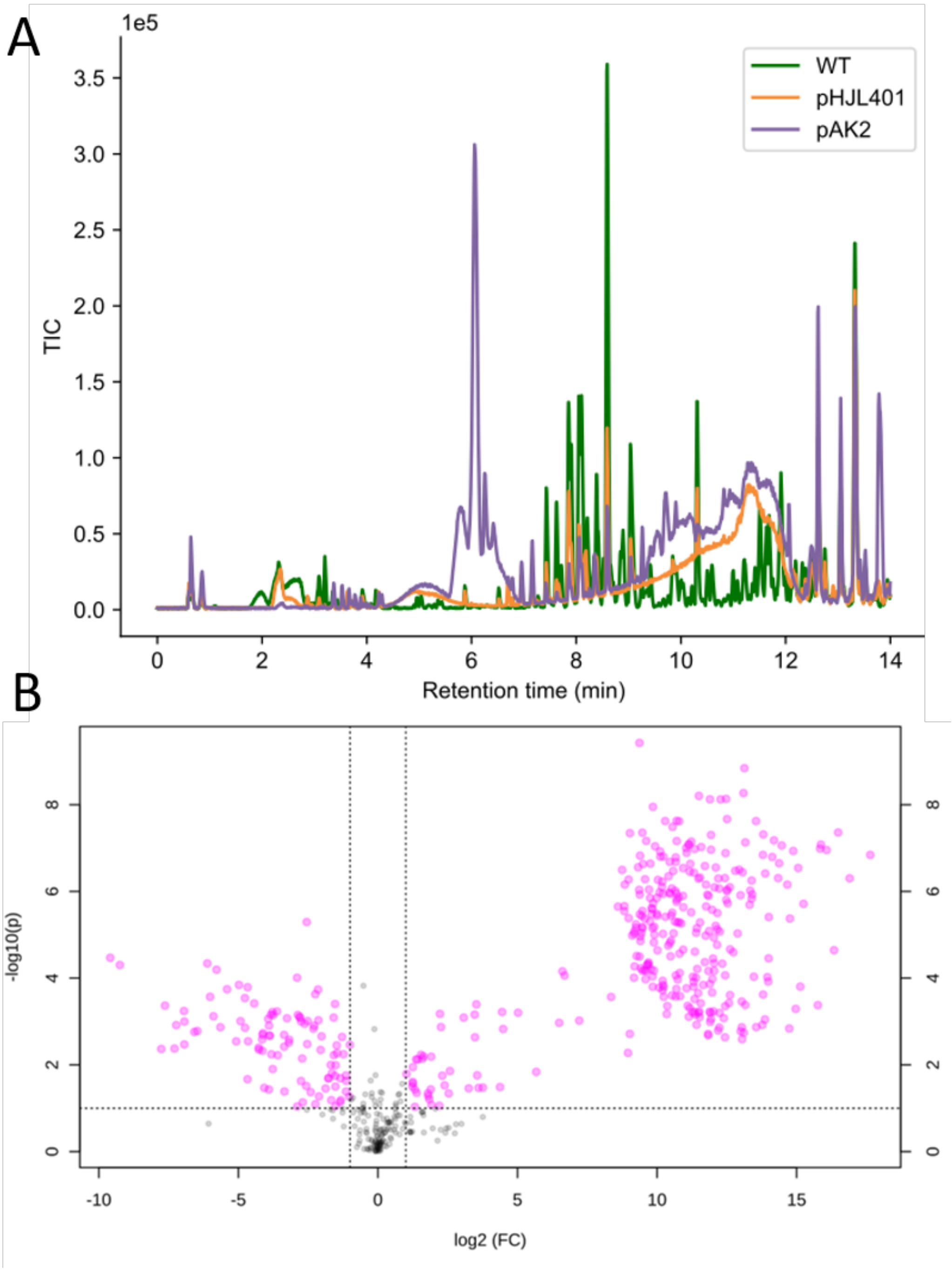

**Figure S13. Transforming the expression construct pAK2 increases the number and TIC of found peaks.** A) Chromatogram of methanol extracts made from *S. pristinaespiralis* harboring nothing (WT) an empty pHJL401 vector (pHJL401), or pAK2 (pHJL401 with *sprR* behind $p_{gap}$). B) Volcano plot comparing extracts of the strain containing pAK2 with the strain containing pHJL401. The two largest peaks were those corresponding to monoisotopic peaks of 2703.245 Da and 2553.260 Da. Several other peaks appeared to be derived from these peaks (table S5).

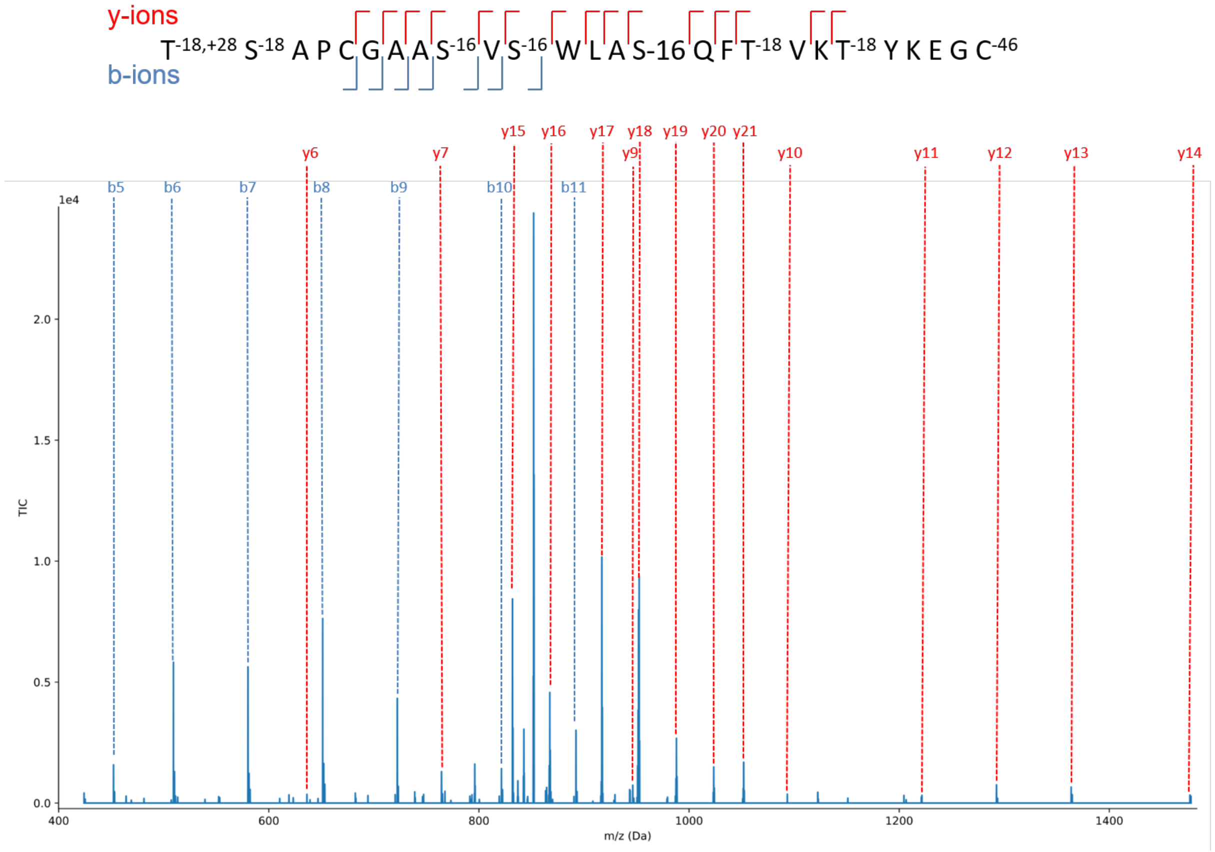

**Figure S14. Fragmentation pattern of the peak corresponding to a monoisotopic mass 2553.260 Da can be matched to the SprA2 precursor.** A large amount of predicted masses of the b- and y-ions can be matched to the predicted modified SprA2 peptide. See table S7 for more details.

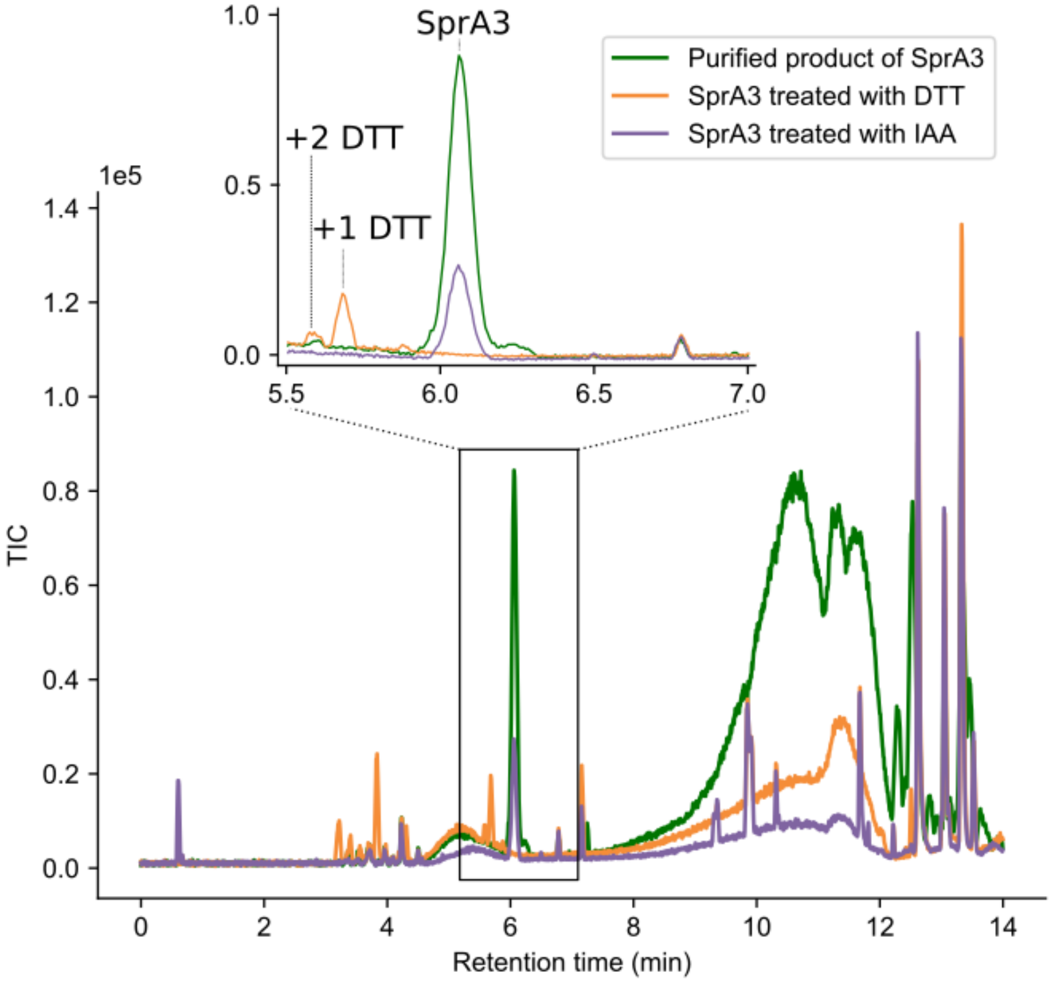

**Figure S15. Labeling experiments with dithiothreithol (DTT) and iodoacetamide (IAA) provide further support for the proposed structure of SprA3.** (Orange) DTT covalently attaches to the alkene group of dehydrated serine and threonine residues. Three of these residues are predicted in the final product of sprA3. DTT labeling shows the presence of up to two adducts, showing indeed that some alkenes are present in the product. Whether the last residue is not correctly predicted or whether adduct formation was hampered due to steric hindrance remains to be determined. (Purple) IAA covalently attaches to free sulfur groups of cysteines. However, the SprA3 peak was unaltered by IAA treatment, despite the presence of three cysteines in the peptide, strongly suggesting that these cysteines are not free.

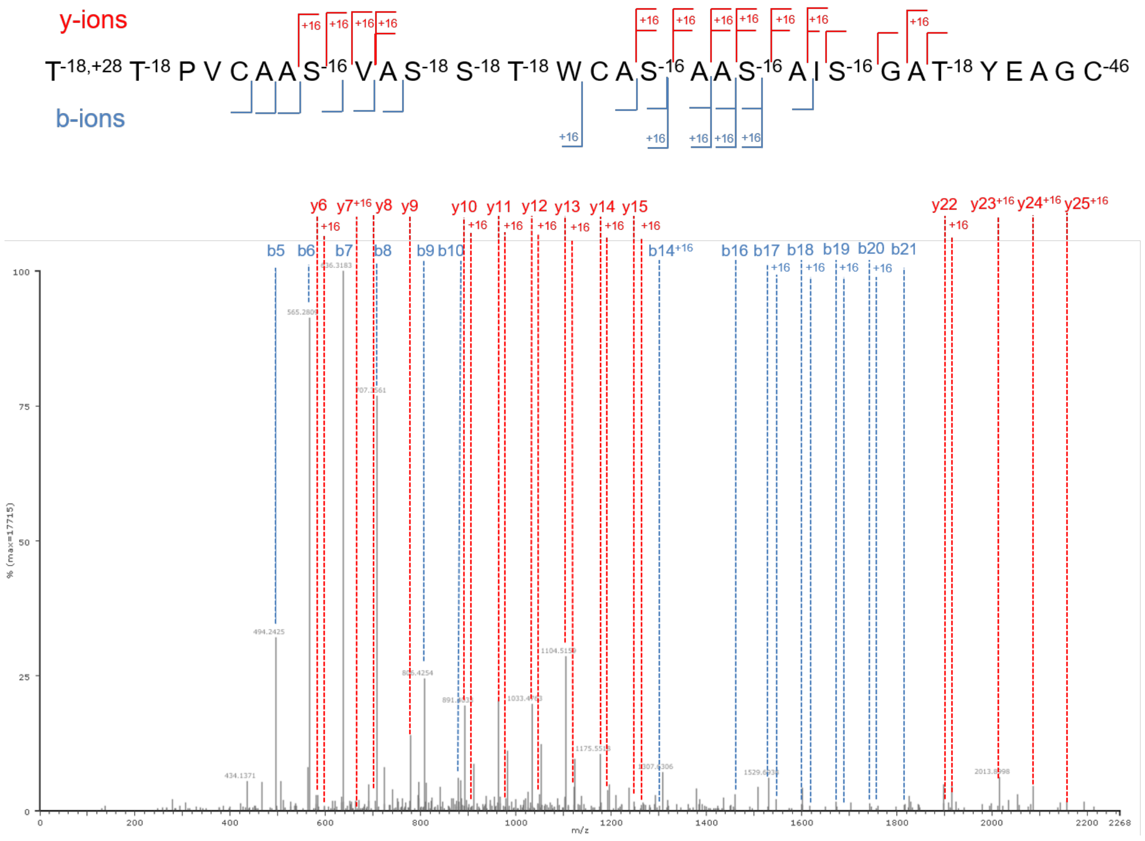

**Figure S16. The fragmentation pattern of the peak corresponding to a monoisotopic mass 2719.245 Da suggests a mixture of unmodified peptides.** Considering that the mass increase was 16 Da compared to the main product, a possible explanation for the mixture of fragments is that the peak detected consisted of a mixture of molecules, each containing a 16 Da modification on a different position. The fragments that are found support this hypothesis, as the N-terminal and C-terminal fragments were found both with and without the 16 Da modification. See table S8 for more details.

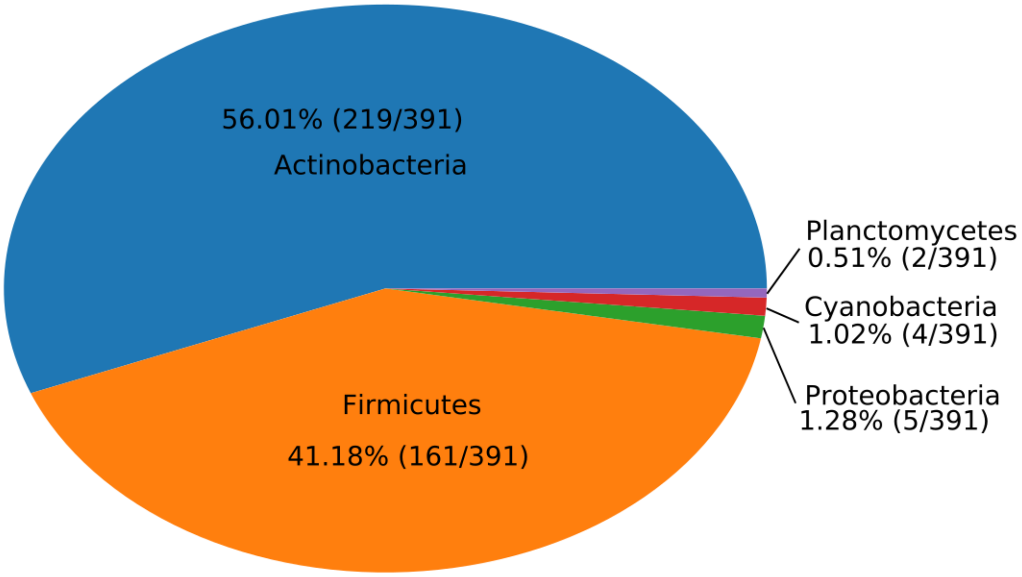

**Figure S17. Homologs of the *sprPT* and *sprH3* gene pair are present outside *Streptomyces*.** Most homologs were found in *Actinobacteria* and *Firmicutes*, although a few additional candidates were found in *Proteobacteria*, *Cyanobacteria* and *Planctomycetes*.

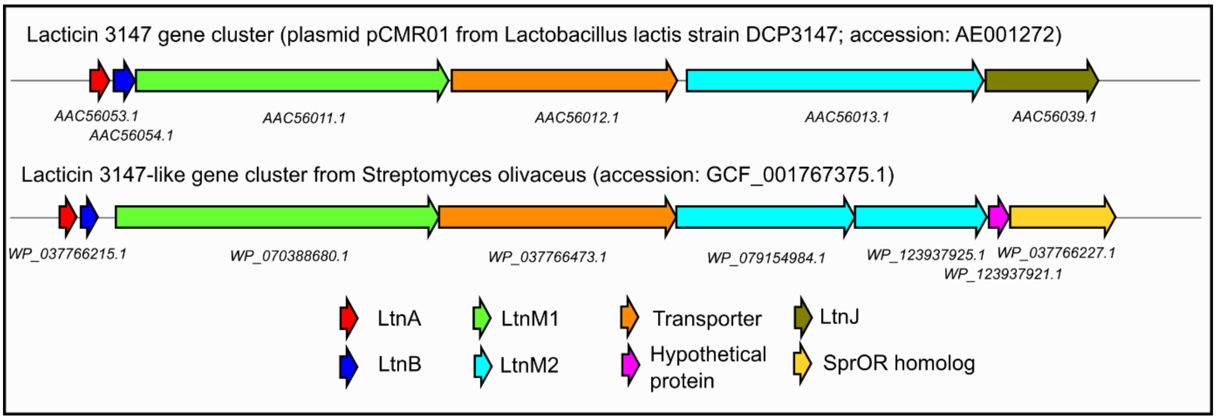

**Figure S18. Comparison of the lacticin 3147-like gene cluster from *Lactococcus lactis* with a homologous cluster from *Streptomyces olivaceus*.** Genes encoding both precursors, both LanM-like modifying enzymes and the transporter are well conserved between the clusters. The gene encoding LtnJ, however, responsible for the reduction in the conversion to alanine and butyric acid, was not conserved. Instead, a homolog to s*prOR* was found, suggesting it may carry out a similar function.

**Table S1. Primers used in this study.**

| Primer name | Primer sequence |
|---|---|
| sprR_F | gatc GAATTC CAT ATGACCGTCAACGACCTGTCC |
| sprR_R | gatc TCTAGA CGCGGCCCACGGATCAGACC |
| spr_LF_F | gatc GAATTC CTCGCGGCCCTCGGCATTCTGG |
| spr_LF_R | gcta TCTAGA GTGGCGTGCGCGGCGTTGG |
| spr_RF_F | gcta TCTAGA CGCCCGGAAACAGGCATGAAGG |
| spr_RF_R | gcta AAGCTT ATGTCGCGGTGGACGACACCC |
| spr_del_check_F | GGGCTACATGCCTACTTTGC |
| spr_del_check_R | GTGCCCTCTGATTCCTTTCC |

**Table S2. Plasmids used in this study.**

| Plasmid name | Description | Reference |
|---|---|---|
| pSET152 | Integrative *E. coli* / *Streptomyces* shuttle vector. | Bierman *et al*[28]. |
| pHJL401 | *E. coli/Streptomyces* shuttle vector with intermediate copy number. | Larson *et al*[29]. |
| pWHM3 | Unstable *E. coli/Streptomyces* shuttle vector with high copy number; used for homologous recombination | Vara *et al*[30]. |
| pUWLCRE | Unstable *E. coli/Streptomyces* shuttle vector containing the Cre recombinase enzyme. behind a constitutive promoter. | Fedorshyn *et al*[31]. |
| pAK1 | pSET152 containing sprR behind GAP promoter from *S. coelicolor* (SCO1947). | This work. |
| pAK2 | pHJL401 containing sprR behind GAP promoter from *S. coelicolor* (SCO1947). | This work. |
| pAK3 | pWHM3 containing regions flanking the SpR gene cluster. | This work. |

**Table S3. Precursor sequences of selected candidate RiPP families.** Serine and threonine resides are marked in green, and cysteine residues are marked in red.

| Family | Genome | Precursor sequence |
|---|---|---|
| Known RiPP markers | Streptomyces sp. NTK 937 | MTENTAPEESPEVEAHSAADDAAQAPEQFHDAAEIICGVYDKEIQV |
| Known RiPP markers | Streptomyces fradiae NKZ-259 | MPSGMPNDPSTTDGLSRRRVLGTAAAAAVPLPARGAEDAEAKSGPW |
| Methylamine utilisation protein MauE | Streptomyces viridosporus T7A | MSRALESLSSRLLGLFVPKVEAAASAQACQCFNECWQCARSACCVNTYCGSINCWRSCPGC |
| Methylamine utilisation protein MauE | Streptomyces sp. CS081A | MARTVGDGSKGCRPSPVSPYGLDQYGDRAASTWGASSATCGVRGEP<br>MVKSLSALAGRAFARVLPQETAAAACACPAGSSSWCSGENLYTRFCCSWNCAAKPTCTVTVVYGAC |
| Methylamine utilisation protein MauE | Streptomyces sp. 2112.3 | MFKKLEAVGSALLERLVPRVDASACGTNCWNDCWQCAHSACKVNTCTGALTCLSGNC |
| ATP-grasp ligases | Streptomyces sp. NRRL F-6491 | MARAARNLLAITASAALSFLLVQGTGAQEERAFLAGSGQGKVINDLGWG |
| ATP-grasp ligases | Streptomyces sp. GSSD-12 | MSSDPSDAAEQGPVGGFITEPLVAAAATTGGCCGEPRSAPEPARSSCCGEPAAEEAPRSCCGEPAAAG<br>MADDMIGSGCCETSGNEDVAEDGTECGCACACCD<br>MSETSLGNMFWNAAQQPPAATAEEPKKASSCCGPKPEAKAPAEQAAAPEKASSCCGPKPAAAAEPEGTPAPKKSSCCG |
| Other | Streptomyces sp. WZ.A104 | MQNVTEKDLFDGYTAYTSAEELGLHDGKEAAPAFSPTIPWAIRATIISARSSQQCAAALGSLAAKTVENKC |
| Other | Streptomyces sp. OspMP-M45 | MTEAGLWEEGDAGRRRPLGVPPENWPVPGGRQGMDGQWSGQSSKTIDHPGGAT |
| Other | Streptomyces avermitilis MA-4680 | MSSLDKPGRKKWSGPEKWQVILAASSLGVAVVALVGQFAQFL |
| Other | Streptomyces koyangensis VK-A60T | MGDLDEEVAAPGPGRWIRPSSTAGYGWTTSCRTSVFPPASPDSCQARETVTWCAWVP |
| Other | Streptomyces sp. ADI95-17 | MNSLSEAGCWCHERLKSCPSECKFRVKDGGAVMKFLFLLKDKMTPEKSLKAYAWYHWY<br>MCEVCRSSRNPGPWGGCCGDGARLGHGWPVSYYETLLCKSQPHEGLDLGASIGEGFEPTPGDLPAGGQSPHKE |
| Other | Streptomyces sp. WAC01280 | MLKGGQLGRFSTNSMNDHREQLGIGPPCLLTFDNAARSSQPSQEAAPCARAES |
| Other | Streptomyces sp. AcE210 | MAESPTPEAVAEQPTEVAQPHRLVLLGACGCGSGCGCGCQSGAPCQCGGCSG |
| Other | Streptomyces punisceus NRRL B-2895 | MRTAAAYASGEPPPVAVVKSHGVAFENRVRYVSPVPSTTHAAASAPGSAEGSAPAATA |
| Other | Streptomyces lydicus ATCC 25470 | MLWKSCARARCGISIPWNSFEFDHGGTGVVPCVPGVCEFPARDGKEEVT<br>MNQGGGEQRGAEVSIRANVGSWLAVRKSPFEAGGSPVSRWEDLPRGVPCPYETGAHQD |

**Table S4. Proteins containing a flavoprotein domain (PF02441) are present in both RiPP and non-RiPP BGCs.** While proteins with this domain are known in RiPP biosynthesis for the decarboxylation of C-terminal cysteines, their presence is not restricted to RiPP BGCs.

| MIBiG BGC ID | BGC class | RiPP class | Protein accession |
|---|---|---|---|
| BGC0000157 | Polyketide | | ABI94381.1 |
| BGC0000158 | Polyketide | | ABV91288.1 |
| BGC0000171 | Polyketide | | CCC21124.1 |
| BGC0000203 | Polyketide | | ADI71473.1 |
| BGC0000203 | Polyketide | | ADI71437.1 |
| BGC0000373 | NRP | | EFG10345.1 |
| BGC0000807 | Saccharide | | ADD45285.1 |
| BGC0000889 | Other | | BAM73626.1 |
| BGC0000932 | Other | | AFO93363.1 |
| BGC0001115 | NRP/Polyketide | | CBK62752.1 |
| BGC0001193 | NRP | | AJI44175.1 |
| BGC0001362 | Other | | AFO93363.1 |
| BGC0001592 | Other | | AVI10267.1 |
| BGC0000508 | RiPP | Lanthipeptide | CAA44255.1 |
| BGC0000514 | RiPP | Lanthipeptide | ABC94905.1 |
| BGC0000527 | RiPP | Lanthipeptide | CAB60260.1 |
| BGC0000529 | RiPP | Lanthipeptide | ADK32557.1 |
| BGC0000530 | RiPP | Lanthipeptide | EMC15126.1 |
| BGC0000531 | RiPP | Lanthipeptide | AAG48568.1 |
| BGC0000533 | RiPP | Lanthipeptide | AAD56146.1 |
| BGC0001618 | RiPP | Lanthipeptide | ARD24448.1 |
| BGC0001669 | RiPP | Lanthipeptide | AVH76813.1 |
| BGC0000582 | RiPP | Linaridin | ADR72965.1 |
| BGC0000583 | RiPP | Linaridin | YP_001827875.1 |
| BGC0000625 | RiPP | Thioamide-containing peptide | BAN83921.1 |
| BGC0001802 | RiPP | Thioamide-containing peptide | ATJ00796.1 |
| BGC0001803 | RiPP | Thioamide-containing peptide | BAN83921.1 |
| BGC0001696 | RiPP | Thioamide-containing peptide | BBC15202.1 |

**Table S5. Peaks unique to strains containing pAK1 appear to be mostly derived from a single mass.**

| Peak m/z | Predicted charge from isotope pattern | Monoisotopic mass (assuming M+H ions) | Description |
|---|---|---|---|
| 707.3534 | 1 | 706.3454 | Fragment of 2703.2349 |
| 868.0891 | 3 | 2601.2433 | |
| 902.0863 | 3 | 2703.2349 | |
| 903.4186 | 3 | 2707.2318 | 2703.2349 + 4 Da (2*$H_2$) |
| 907.4167 | 3 | 2719.2261 | 2703.2349 + 16 Da (O) |
| 908.7487 | 3 | 2723.2221 | 2703.2349 + 20 Da (O + 2*$H_2$) |
| 914.9003 | NA | NA | |
| 918.0807 | 3 | 2751.2181 | 2703.2349 + 48 Da (3*O) |

**Table S6. Many detected masses from strains containing the expression construct pAK2 appear to be derived from two masses.** The two base masses were also the most abundant, making it likely these form final products, while the other masses may be incompletely processed products.

| Description | Calculated m/z M+3H (Da) | Observed m/z pAK2 | delta ppm |
|---|---|---|---|
| **Most abundant mass #1** | 902.088 | 902.085 | 3.3 |
| **+ oxygen** | 907.420 | 907.417 | 2.8 |
| | 907.420 | 907.417 | 2.4 |
| **+2 oxygen** | 912.751 | 912.748 | 3.6 |
| | 912.751 | 912.750 | 1.3 |
| **+3 oxygen** | 918.083 | 918.080 | 2.6 |
| | 918.083 | 918.081 | 1.6 |
| | 918.083 | 918.082 | 1.2 |
| **+4 oxygen** | 923.414 | 923.412 | 3.1 |
| **+ methyl** | 906.760 | 906.757 | 3.5 |
| **- methyl** | 897.416 | 897.415 | 0.8 |
| **- 2 methyl** | 892.744 | | |
| **Most abundant mass #2** | 852.097 | 852.095 | 2.3 |
| **+ oxygen** | 852.376 | 857.426 | 0.9 |
| **+2 oxygen** | 862.758 | 862.758 | 0.0 |
| **+3 oxygen** | 868.090 | 868.090 | 0.6 |
| | 868.090 | 868.089 | 0.7 |
| **- 2 methyl** | 842.751 | 842.747 | 4.2 |

**Table S7. Observed masses for fragments of a mass of 2703.235 Da can be matched to the SprA3 precursors.** See Figure 3 (main text) for more details.

| Ion | Calculated mass | Observed mass | Δppm | Ion | Calculated mass | Observed mass | Δppm |
|---|---|---|---|---|---|---|---|
| b1 | 112.08 | | | y30 | 2593.17 | | |
| b2 | 195.11 | | | y29 | 2510.13 | | |
| b3 | 292.17 | | | y28 | 2413.08 | | |
| b4 | 391.23 | | | y27 | 2314.01 | | |
| b5 | 494.24 | 494.2412 | 5.5 | y26 | 2211.00 | | |
| b6 | 565.28 | 565.2805 | 0.9 | y25 | 2139.97 | | |
| b7 | 636.32 | 636.3193 | 1.9 | y24 | 2068.93 | | |
| b8 | 707.36 | 707.3570 | 2.5 | y23 | 1997.89 | | |
| b9 | 806.42 | 806.4266 | 3.7 | y22 | 1898.82 | | |
| b10 | 877.46 | 877.4478 | 14.8 | y21 | 1827.79 | | |
| b11 | 946.48 | | | y20 | 1758.77 | | |
| b12 | 1015.50 | | | y19 | 1689.74 | | |
| b13 | 1098.54 | | | y18 | 1606.71 | | |
| b14 | 1284.62 | | | y17 | 1420.63 | | |
| b15 | 1387.63 | | | y16 | 1317.62 | 1317.608 | 8.1 |
| b16 | 1458.67 | | | y15 | 1246.58 | 1246.593 | 9.4 |
| b17 | 1529.70 | 1529.7040 | 0.3 | y14 | 1175.54 | 1175.557 | 11.3 |
| b18 | 1600.74 | | | y13 | 1104.51 | | |
| b19 | 1671.78 | | | y12 | 1033.47 | | |
| b20 | 1742.81 | | | y11 | 962.43 | 962.436 | 3.4 |
| b21 | 1813.85 | | | y10 | 891.40 | | |
| b22 | 1926.94 | | | y9 | 778.31 | | |
| b23 | 1997.97 | | | y8 | 707.27 | 707.277 | 3.9 |
| b24 | 2054.99 | | | y7 | 650.25 | 650.256 | 4.9 |
| b25 | 2126.03 | | | y6 | 579.22 | 579.211 | 7.9 |
| b26 | 2209.07 | | | y5 | 496.18 | | |
| b27 | 2372.13 | | | y4 | 333.12 | | |
| b28 | 2501.17 | | | y3 | 204.07 | | |
| b29 | 2572.21 | | | y2 | 133.04 | | |
| b30 | 2629.23 | | | y1 | 76.01 | | |

**Table S8. Observed masses for fragments of a peak corresponding to a monoisotopic mass of 2553.260 Da can be matched to the SprA2 precursor.** See Figure S10 for more details.

| Ion | Calculated m/z (Da) | Observed m/z | Δppm | Ion | Calculated m/z (Da) | Calculated m/z (doubly charged) | Observed m/z | Δppm |
|---|---|---|---|---|---|---|---|---|
| b1 | 112.0771 | | | y1 | 76.02193 | | | |
| b2 | 181.0986 | | | y2 | 133.0434 | | | |
| b3 | 252.1357 | | | y3 | 262.086 | | | |
| b4 | 349.1885 | | | y4 | 390.1809 | | | |
| b5 | 452.1977 | 452.1976 | 0.1 | y5 | 553.2443 | | | |
| b6 | 509.2191 | 509.219 | 0.2 | y6 | 636.2814 | | 636.2841 | 4.3 |
| b7 | 580.2562 | 580.2552 | 1.8 | y7 | 764.3764 | | 764.3755 | 1.1 |
| b8 | 651.2933 | 651.2933 | 0.0 | y8 | 863.4448 | | | |
| b9 | 722.3304 | 722.3316 | 1.6 | y9 | 946.4819 | | 946.4829 | 1.1 |
| b10 | 821.3989 | 821.3968 | 2.5 | y10 | 1093.55 | | 1093.554 | 3.5 |
| b11 | 892.436 | 892.4338 | 2.4 | y11 | 1221.609 | | 1221.606 | 2.1 |
| b12 | 1078.515 | | | y12 | 1292.646 | | 1292.651 | 3.9 |
| b13 | 1191.599 | | | y13 | 1363.683 | | 1363.672 | 8.1 |
| b14 | 1262.636 | | | y14 | 1476.767 | | 1476.769 | 1.0 |
| b15 | 1333.674 | | | y15 | 1662.846 | 831.9272 | 831.9276 | 0.4 |
| b16 | 1461.732 | | | y16 | 1733.884 | 867.4458 | 867.4469 | 1.3 |
| b17 | 1608.801 | | | y17 | 1832.952 | 916.98 | 916.9802 | 0.2 |
| b18 | 1691.838 | | | y18 | 1903.989 | 952.4986 | 952.4948 | 3.9 |
| b19 | 1790.906 | | | y19 | 1975.026 | 988.0171 | 988.0172 | 0.1 |
| b20 | 1919.001 | | | y20 | 2046.063 | 1023.536 | 1023.54 | 4.0 |
| b21 | 2002.038 | | | y21 | 2103.085 | 1052.046 | 1052.043 | 3.6 |
| b22 | 2165.101 | | | y22 | 2206.094 | | | |
| b23 | 2293.196 | | | y23 | 2303.147 | | | |
| b24 | 2422.239 | | | y24 | 2374.184 | | | |
| b25 | 2479.26 | | | y25 | 2443.205 | | | |

**Table S9. Fragments of a peak with corresponding to a monoisotopic mass of 2719.245 Da matched to the b- and y-ions.** Fragments could both be matched when considering the base product, or the same product incremented by the mass of a single oxygen atom (~16 Da), relating to a single unmodified serine residue. Since fragments for the same b-ion are observed both with and without this mass difference, this peak likely represents a mixture of different compounds, each with an unaltered serine in a different position.

| Ion | Calculated mass (Da) | Observed mass (Da) | Δppm | Calculated mass + oxygen (Da) | Observed mass (Da) | Δppm |
|---|---|---|---|---|---|---|
| b1 | 112.08 | | | 128.07133 | | |
| b2 | 195.11 | | | 211.10845 | | |
| b3 | 292.17 | | | 308.16121 | | |
| b4 | 391.23 | | | 407.22962 | | |
| b5 | 494.24 | 494.2425 | 2.83 | 510.23881 | | |
| b6 | 565.28 | 565.2809 | 0.19 | 581.27592 | | |
| b7 | 636.32 | 636.3183 | 0.28 | 652.31303 | | |
| b8 | 707.36 | 707.3561 | 1.22 | 723.35015 | | |
| b9 | 806.42 | 806.4254 | 2.17 | 822.41856 | | |
| b10 | 877.46 | 877.4634 | 3.01 | 893.45567 | | |
| b11 | 946.48 | | | 962.47714 | | |
| b12 | 1015.50 | | | 1031.49861 | | |
| b13 | 1098.54 | | | 1114.53573 | | |
| b14 | 1284.62 | | | 1300.61504 | 1300.608 | 5.41 |
| b15 | 1387.63 | | | 1403.62423 | | |
| b16 | 1458.67 | 1458.6644 | 1.39 | 1474.66134 | | |
| b17 | 1529.70 | 1529.6938 | 6.37 | 1545.69846 | 1545.7004 | 1.26 |
| b18 | 1600.74 | 1600.7327 | 4.97 | 1616.73557 | 1616.741 | 3.36 |
| b19 | 1671.78 | 1671.7671 | 6.38 | 1687.77268 | 1687.751 | 12.85 |
| b20 | 1742.81 | | | 1758.80979 | 1758.7939 | 9.03 |
| b21 | 1813.85 | 1813.8336 | 10.14 | 1829.84691 | | |
| b22 | 1926.94 | | | 1942.93097 | | |
| b23 | 1997.97 | | | 2013.96809 | | |
| b24 | 2054.99 | | | 2070.98955 | | |
| b25 | 2126.03 | | | 2142.02666 | | |
| b26 | 2209.07 | | | 2225.06378 | | |
| b27 | 2372.13 | | | 2388.12711 | | |
| b28 | 2501.17 | | | 2517.1697 | | |
| b29 | 2572.21 | | | 2588.20681 | | |
| b30 | 2629.23 | | | 2645.22827 | | |

**Table S9 (continued). Fragments of a peak with 2719.245 dalton matched to the b- and y-ions.**

| Ion | Calculated mass (Da) | Observed mass (Da) | Δppm | Calculated mass + oxygen (Da) | Observed mass (Da) | Δppm |
|---|---|---|---|---|---|---|
| y30 | 2593.17 | | | 2609.17 | | |
| y29 | 2510.13 | | | 2526.13 | | |
| y28 | 2413.08 | | | 2429.08 | | |
| y27 | 2314.01 | | | 2330.01 | | |
| y26 | 2211.00 | | | 2227 | | |
| y25 | 2139.97 | 579.219 | 5.7 | 2155.96 | 2155.98 | 6.6 |
| y24 | 2068.93 | 650.265 | 18.0 | 2084.92 | 2084.94 | 7.1 |
| y23 | 1997.89 | 707.278 | 5.6 | 2013.89 | 2013.9 | 6.2 |
| y22 | 1898.82 | 778.319 | 9.2 | 1914.82 | 1914.79 | 14.6 |
| y21 | 1827.79 | 891.403 | 8.6 | 1843.78 | | |
| y20 | 1758.77 | 962.44 | 7.6 | 1774.76 | | |
| y19 | 1689.74 | 1033.48 | 6.2 | 1705.74 | | |
| y18 | 1606.71 | 1104.52 | 8.1 | 1622.7 | | |
| y17 | 1420.63 | 1175.55 | 6.6 | 1436.62 | | |
| y16 | 1317.62 | 1246.59 | 6.2 | 1333.61 | 1333.61 | 5.6 |
| y15 | 1246.58 | 1317.64 | 15.1 | 1262.58 | 1262.59 | 14.1 |
| y14 | 1175.54 | | | 1191.54 | 1191.55 | 10.6 |
| y13 | 1104.51 | | | 1120.5 | 1120.52 | 12.0 |
| y12 | 1033.47 | 1689.71 | 18.4 | 1049.46 | 1049.46 | 1.0 |
| y11 | 962.43 | 1758.79 | 16.2 | 978.428 | 978.43 | 2.2 |
| y10 | 891.40 | | | 907.391 | 907.379 | 13.3 |
| y9 | 778.31 | 1898.83 | 1.0 | 794.306 | | |
| y8 | 707.27 | | | 723.269 | | |
| y7 | 650.25 | | | 666.248 | 666.251 | 5.1 |
| y6 | 579.22 | | | 595.211 | 595.218 | 11.3 |
| y5 | 496.18 | | | 512.174 | | |
| y4 | 333.12 | | | 349.11 | | |
| y3 | 204.07 | | | 220.068 | | |
| y2 | 133.04 | | | 149.031 | | |
| y1 | 76.01 | | | 92.0092 | | |

**Table S10. Cysteines linked to serine and threonine residues are detected after acidic hydrolysis of SprA3.** Below are the predicted amino acids, and their detected masses. Most of the amino acids can be detected in the chromatogram, including the cysteines linked to dehydrated serine and threonine residues. The mass of the predicted decarboxylated cysteine linked to a dehydrated threonine residue was not detected, nor were the dehydrated serine and threonine residue. However, given that these groups contain alkenes, which easily react under acidic conditions, these groups may have been degraded.

| Amino acid | Calculated mass | M+H | Detected m/z | Δppm |
|---|---|---|---|---|
| Glycine | 75.032 | 76.040 | NA | NA |
| Serine $^{-18}$ | 87.032 | 88.040 | NA | NA |
| Alanine/Serine$^{-16}$ | 89.048 | 90.056 | 90.055 | 13.0 |
| Threonine $^{-18}$ | 101.048 | 102.056 | NA | NA |
| Proline | 115.064 | 116.072 | 116.071 | 8.9 |
| Valine | 117.079 | 118.087 | 118.086 | 9.3 |
| Isoleucine | 131.095 | 132.103 | 132.102 | 7.9 |
| Glutamate | 147.054 | 148.062 | 148.060 | 7.1 |
| Decarboxylated cysteine – threonine | 176.062 | 177.070 | NA | NA |
| Tyrosine | 181.074 | 182.082 | 182.081 | 5.5 |
| Tryptophan | 204.090 | 205.098 | NA | NA |
| Cysteine – Serine | 208.052 | 209.060 | 209.059 | 2.0 |
| Cysteine – Threonine (twice methylated) | 250.100 | 251.108 | 251.106 | 5.5 |

**Table S11. Homologs of the genes *lanJ$_A$*, *sprF1, sprF2 and sprOR* are found associated with both known lanthipeptide BGCs and close to the *sprPT/sprH3* gene pair.** Homology was determined at a cutoff of 30% amino acid identity of the gene products. Within *Streptomyces* genomes, all homologs were found within the analyzed 1,295 genomes. It was then checked whether these homologs overlapped with an antiSMASH-detected lanthipeptide BGC, or were within 15 genes of the *sprPT/sprH3* gene pair. *sprOR* homologs were found within canonical lanthipeptide BGCs as well as associated with the *spriPT/sprH3* gene pair, suggesting its association with lanthipeptide BGCs.

For non-*Streptomyces* genomes, the *sprPT/sprH3* gene pair was first detected, and homologs of the given queries were found within the 15 surrounding genes. Homologs of *lanJ$_A$* and *sprF1* are often found associated with *sprPT/sprH3* gene pair, suggesting they are involved in lanthipeptide biosynthesis.

| ***Streptomyces* genomes** | | | | |
|---|---|---|---|---|
| **Query** | **Overlap with lanthipeptide BGC** | ***sprPT/sprH3* gene pair** | **Overlap with both** | **Overlap with neither** |
| ***lanjA*** | 0 | 0 | 0 | 5 |
| ***sprOR*** | 124 | 137 | 2 | 199 |
| ***sprF1*** | 0 | 124 | 2 | 16 |
| ***sprF2*** | 13 | 135 | 2 | 348 |
| **Non-*Streptomyces* gene clusters** | | | | |
| **Query** | **Overlap with lanthipeptide BGC** | ***sprPT/sprH3* gene pair** | **Overlap with both** | **Overlap with neither** |
| ***lanjA*** | 0 | 40 | 0 | 0 |
| ***sprOR*** | 0 | 108 | 0 | 0 |
| ***sprF1*** | 0 | 111 | 0 | 0 |
| ***sprF2*** | 0 | 146 | 0 | 0 |
